# Structure and dynamics determine G protein coupling specificity at a class A GPCR Teaser: Structure and dynamics studies reveal a mechanism for GPCR signaling bias and G-protein coupling specificity

**DOI:** 10.1101/2024.03.28.587240

**Authors:** Marina Casiraghi, Haoqing Wang, Patrick Brennan, Chris Habrian, Harald Hübner, Maximilian F. Schmidt, Luis Maul, Biswaranjan Pani, Sherif M.F.M. Bahriz, Bing Xu, Nico Staffen, Tufa E. Assafa, Bohan Chen, Elizabeth White, Roger K. Sunahara, Asuka Inoue, Yang K. Xiang, Robert J. Lefkowitz, Ehud Y. Isacoff, Nathaniel Nucci, Peter Gmeiner, Michael T. Lerch, Brian K. Kobilka

## Abstract

G protein coupled receptors (GPCRs) exhibit varying degrees of selectivity for different G protein isoforms. Despite the abundant structures of GPCR-G protein complexes, little is known about the mechanism of G protein coupling specificity. The β_2_-adrenergic receptor is an example of GPCR with high selectivity for Gαs, the stimulatory G protein for adenylyl cyclase, and much weaker for the Gαi family of G proteins inhibiting adenylyl cyclase. By developing a Gαi-biased agonist (LM189), we provide structural and biophysical evidence supporting that distinct conformations at ICL2 and TM6 are required for coupling of the different G protein subtypes Gαs and Gαi. These results deepen our understanding of G protein specificity and bias and can accelerate the design of ligands that select for preferred signaling pathways.

## Introduction

There are over 800 members of the G protein coupled receptor (GPCR) superfamily (*1*), yet they couple with varying efficacy to only four G protein subfamilies (Gαs, Gαi/o, Gαq/11, Gα12/13) to activate distinct downstream signaling cascades (*2*). In recent years, the structures of over 400 GPCR-G protein complexes with different G protein subtypes have been reported (*3–9*). However, the molecular determinants of GPCR-G protein coupling specificity remain largely unknown (*10–14*). In addition, mutagenesis and phylogenetic analysis have found no correlation between sequence and coupling selectivity (*15*, *16*). Biophysical investigations have shown that GPCRs are inherently flexible, existing in an equilibrium of multiple conformations (*17–20*). Depending on their efficacy, ligands can shift this equilibrium towards specific states, facilitating the coupling of signaling partners. However, partner-specific states are likely transient, low-probability conformations, that cannot be trapped by structural methods such as X-ray crystallography or cryo-EM. Therefore, additional biophysical studies are needed to complement the information provided by structures and to delineate the transient yet important conformational states stabilized in the absence of bound G proteins(*21–24*).

A deeper understanding of the mechanism at the basis of G protein specificity is essential for the development of drugs that preferentially activate a single G protein subtype. This could reduce the potential adverse effects associated with the activation of multiple G proteins isoforms. In this context, the development of biased ligands that preferentially activate a single G protein subtype is highly desirable as a tool to better characterize GPCR signaling and for therapeutic purposes. However, our understanding of the molecular determinants underlying biased signaling is still fragmentary, suggesting the need for a more detailed description of the conformational states adopted by receptors bound to biased ligands (*21–24*).

We chose the β_2_-adrenergic receptor (β_2_AR) as a prototypical class A GPCR to investigate G protein specificity and biased signaling. β_2_AR and β_1_-adrenergic receptors (β_1_AR) are GPCRs expressed in cardiac myocytes and play essential roles in the regulation of cardiac function by the sympathetic nervous system. β_1_AR and β_2_AR primarily couple to the stimulatory G protein for adenylyl cyclase Gαs, to increase heart rate and contractility (*25*). β_2_AR also binds to the Gαi subtype, the inhibitory G protein for adenylyl cyclase; activation of Gαi by the β_2_AR can counteract the effects of Gαs activation on heart rate and contractility (*26*). β_2_AR signaling through Gαi can also lead to activation of MAPK/ERK and PI-3K pathways. Chronic stimulation of the Gαs pathway leads to pathologic changes in the heart including myocyte apoptosis, that ultimately leads to congestive heart failure. In contrast, β_2_AR activation of Gαi has a cardioprotective effect by activating the PI3K-Akt signaling cascade (*27*, *28*). However, increased Gαi signaling by the β_2_AR has also been linked to the acceleration of pathologic changes in non-ischemic models of heart failure (*29*), underlying the importance of understanding the molecular basis of the promiscuous signaling through both Gαs and Gαi.

Gαs recruitment to the β_2_AR has been characterized by structural and spectroscopic methods, including NMR and DEER spectroscopy (*4*, *17*, *18*, *23*, *30–33*). In contrast, limited information is available regarding the binding of Gαi to the β_2_AR. Receptor phosphorylation has been proposed to play a role in Gαi recruitment at the β_2_AR (*34*). However, previous biochemical studies showed that in vitro receptor phosphorylation with PKA failed to enhance Gαi coupling to the receptor (*35*). In this study, in vitro receptor phosphorylation with PKA failed to enhance G-protein recruitment to the β_2_AR with all the Gαi-protein subtypes tested (Gαi_1_, Gαi_2_, Gαi_3_). Moreover, previous investigations in neonatal cardiac myocytes showed a biphasic coupling of the β_2_AR to Gs followed by Gi. In this experiments, Gi coupling was not affected by the PKA inhibitor PKI (*36*).

In this work, we identified a Gαi-biased agonist (LM189) for the β_2_AR and investigated the mechanism for its signaling bias using a combination of structural and biophysical methods. We observed that relative to non-biased agonists, LM189 stabilizes a distinct conformation in TM6 and increases the dynamics of ICL2, explaining the preferential Gαi bias mediated by LM189 at this receptor system.

## Results

### Development of the G*α*i-Biased Ligand LM189

Structures of GPCRs coupled to the inhibitory G protein Gi show a smaller outward movement of TM6 compared to Gs and Gq/11 complexes (*3–7*, *9*, *37*). This has been attributed to the smaller size of the C-terminus of the alpha-5 helix of Gαi compared to the bulkier alpha5 of the Gαs and Gαq subtypes (*9*, *13*). A smaller outward displacement of TM6 is also observed in the crystal structure of the β_2_AR bound to the partial agonist salmeterol (*38*). To screen for ligands that could stabilize the β_2_AR-Gi complex for structure determination, we compared the effect of salmeterol with other β_2_AR agonists for the coupling to Gs and Gi using the GTPase-Glo™ Assay (*32*) (**Fig. 1, A-B**) (Note we will use Gs and Gi, instead of Gαs and Gαi, when referring to the heterotrimer). Salmeterol is a subtype-selective β_2_AR partial agonist for Gs activation relative to epinephrine, the endogenous β_2_AR hormone also known as adrenaline (**Fig. 1A, C**). Surprisingly, the GTPase assay showed that salmeterol is more efficacious than epinephrine for β_2_AR coupling to Gi (**Fig. 1B**). Salmeterol is a long-acting β_2_AR agonist (LABA) composed of a saligenin ethanolamine pharmacophore and an aryloxy alkyl tail (*38*, *39*) (**Fig. 1C**). The high subtype selectivity for β_2_AR is mediated by binding of the salmeterol tail in the receptor extracellular vestibule (also known as the exosite) (*38*). Interestingly, salbutamol, another β_2_AR partial agonist that shares the same saligenin ring as salmeterol but lacks the aryloxy alkyl tail, did not display high efficacy for Gi (**Fig. 1A-C**). This suggests that the increased Gαi efficacy of salmeterol is, in part, mediated by its tail (**Fig. 1C**).

**Fig. 1.**
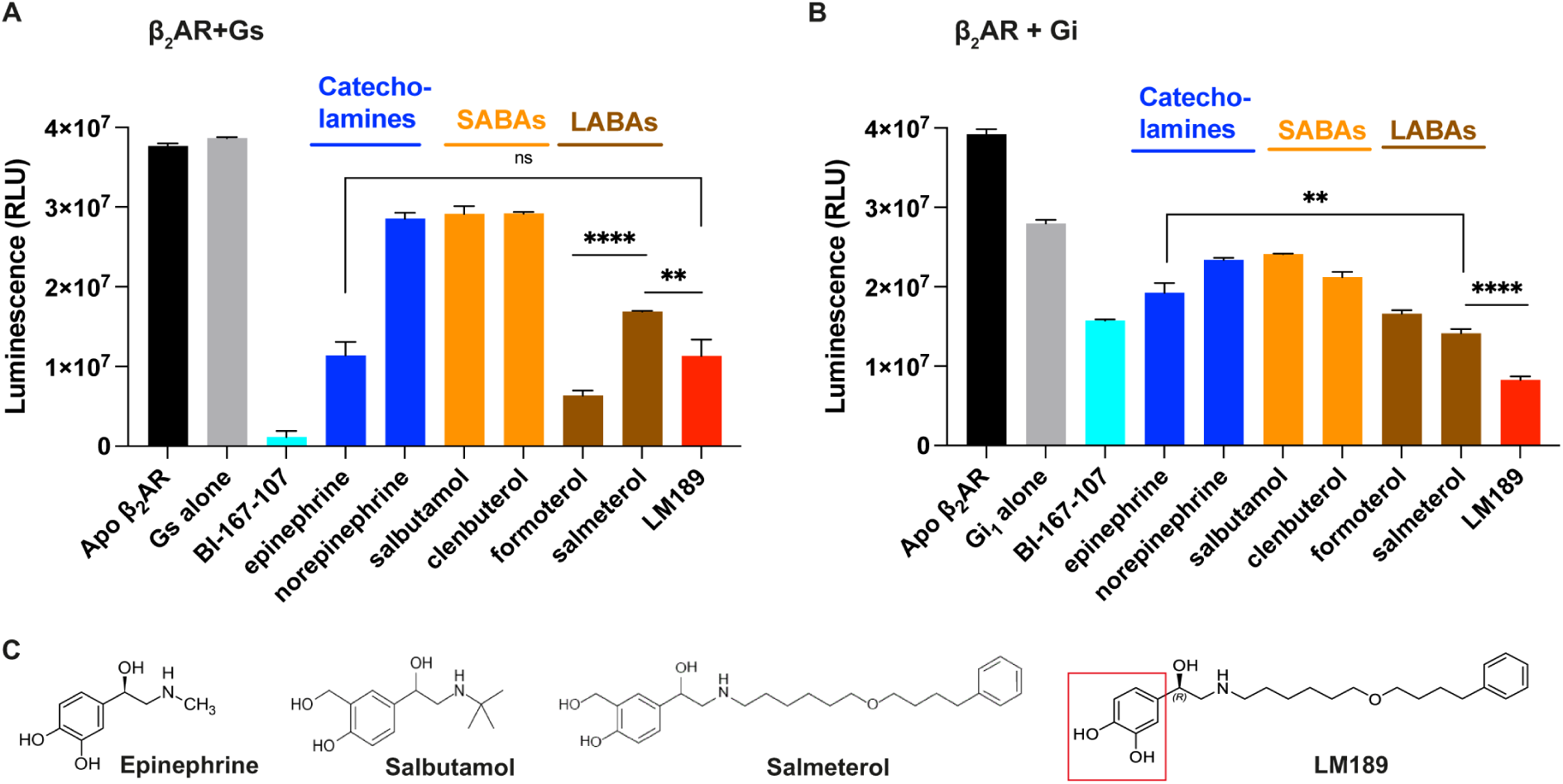
Ligand efficacy at the *β*_2_AR. **(A-B)** Luminescence GTPase-Glo assay. Ligand efficacy reflects the ability of the ligand-bound receptor to promote turnover of the G protein cycle. The reaction starts when β_2_AR, bound to different ligands, is incubated with Gs **(A)** or Gi **(B)**. At the end of the reaction, lower bars correspond to higher GTPase activity. **(A)** Salmeterol is a subtype-selective β_2_AR partial agonist for Gs activation relative to other agonist such as epinephrine and formoterol (\*\*\*\**P*<0.0001). LM189 is as efficacious as epinephrine at Gs turnover (ns, *P*=0.96). In **(B)**, salmeterol is more efficacious than epinephrine for β_2_AR coupling to Gi (\*\**P*=0.003). LM189 is more efficacious than epinephrine and salmeterol at coupling to Gi (\*\*\*\**P*=0.0001). Experiments were performed as biological triplicates and results were plotted using GrapPad Prism. P values were calculated using the unpaired t test analysis on GraphPad Prism, assuming Gaussian distributions. Ns=(*P*>0.05), * (*P*≤0.05), ** (*P*≤0.01), *** (*P*≤0.001), **** (*P*≤0.0001). Data are represented as the mean ± s.d. **(C)** Structures of β_2_AR ligands. From the left, epinephrine (adrenaline) is the endogenous catecholamine neurotransmitter. Salbutamol is a partial agonist belonging to the short acting β_2_AR agonists (SABAs). Salmeterol is a long-acting β_2_AR partial agonist (LABA), which exhibits a long duration of action and is used in the chronic management of asthma. Salmeterol has the same saligenin head group as salbutamol. LM189 was developed by replacing the saligenin moiety of salmeterol with a catechol group (red square).

To better understand the increased efficacy of salmeterol for the coupling of β_2_AR to Gi, we sought to determine the Cryo-EM structure of the complex. Despite our efforts, we could not obtain the structure of the salmeterol-bound β_2_AR-Gi complex. We identified conditions that led to a biochemically stable interaction, however the complex dissociated upon sample vitrification. To further enhance the ligand efficacy for Gαi activation, we designed alternative ligands starting from the salmeterol scaffold. One of the ligands tested, named LM189, proved to be more efficacious than salmeterol at Gi turnover (**Fig. 1 B, C**). LM189 shares the tail region of salmeterol, while the saligenin moiety has been replaced by the catechol group, similar to epinephrine (**Fig. 1C**). LM189 is more efficacious than epinephrine and salmeterol at coupling to Gi (**Fig. 1B**) and as efficacious as epinephrine at Gs turnover (**Fig. 1A**). LM189 is equally efficacious for the Gi_1_, Gi_2_ and Gi_3_ subtypes (**Fig. S1A, B**). For further experiments, we decided to focus on the Gi_1_ subtype that will henceforth be referred to as Gi. Similar results in the GTPase assay were obtained with receptor reconstituted in HDLs (**Fig. S1C**).

To quantify the degree of bias of LM189, we performed nano-BRET experiments between β_2_AR-Rluc and mini-Gs-venus and mini-Gs/i-venus (**Fig. S1D**). We obtained dose-response curves for Gs and Gi activation in the presence of epinephrine, the endogenous β_2_AR ligand that we chose as reference, formoterol and LM189. We used the Operational Model equation (*40*, *41*) in Graphpad Prism to fit the data and determine the LogR values (equivalent to log(t/KA) ratios) (**Fig. S1D**). We then calculated ΔLog(τ/KA) ratios, SEM and relative effectiveness (RE) considering epinephrine as the reference ligand (*40*, *41*)(**Fig. S1D**). ΔΔLog(τ/KA) ratios and bias factors (BF) for LM189 activation of Gi were also determined. Based on our calculations, formoterol is weakly biased at Gi (BF=3.7, **Fig. S1D**), while LM189 shows significant bias towards Gi (BF=24, **Fig. S1D**).

We further characterized LM189-mediated beta-arrestin recruitment profile, that we found similar to epinephrine (**Fig. S1E**). Based on radioligand binding measurements, LM189 shows very high affinity for the β_2_AR (*Ki*=0.063 nM) but can also bind to the β_1_AR with lower affinity (*Ki*= 28 nM) (**Fig. S1F, Table S1**).

Collectively, our experimental data support the higher efficacy of LM189 over balanced ligands at Gi recruitment to the β_2_AR.

### Structure of the LM189-Bound *β*_2_AR-Gi Complex

To determine the binding mode of LM189 and the structural determinants of Gi coupling, we obtained the structure of the β_2_AR-Gi complex by cryo-EM to a global resolution of 2.9 Å (**Fig. 2A-B, Fig. S2, S3A-F**). The overall structure of the LM189-β_2_AR-Gi complex is similar to the previously determined BI-167107-β_2_AR-Gs crystal structure (*4*) (**Fig. 2B**). The position of the alpha5 helix in the β_2_AR-Gi structure closely resembles the β_2_AR-Gs complex, in contrast to other GPCR-Gi complexes such as the µOR-Gi (*3–5*) (**Fig. 2C**). Conversely, TM6 of the receptor in the β_2_AR-Gi complex displays a larger outward movement, more similar to the one observed in the β_2_AR-Gs structure rather than the smaller TM6 opening reported for other GPCR-Gi structures (*3–5*) (**Fig. 2C**). Similar to the β_2_AR-Gs structure, Phe 139 of β_2_AR ICL2 inserts into the hydrophobic pocket of Gi, formed by Phe 191 and Leu 194 of the beta1 strand, Phe 336, Thr 340, Ile 343, Ile 344 of the alpha5 C-terminus (**Fig. 2D**). However, compared to the Gs structure, engagement of Gi to the β_2_AR is mediated by far fewer contacts between the G protein alpha5 and ICL2, TM3 and TM5 of the receptor, accounting for the diminished efficacy and less stable interaction of the β_2_AR with Gi (**Fig. 2D**).

**Fig. 2.**
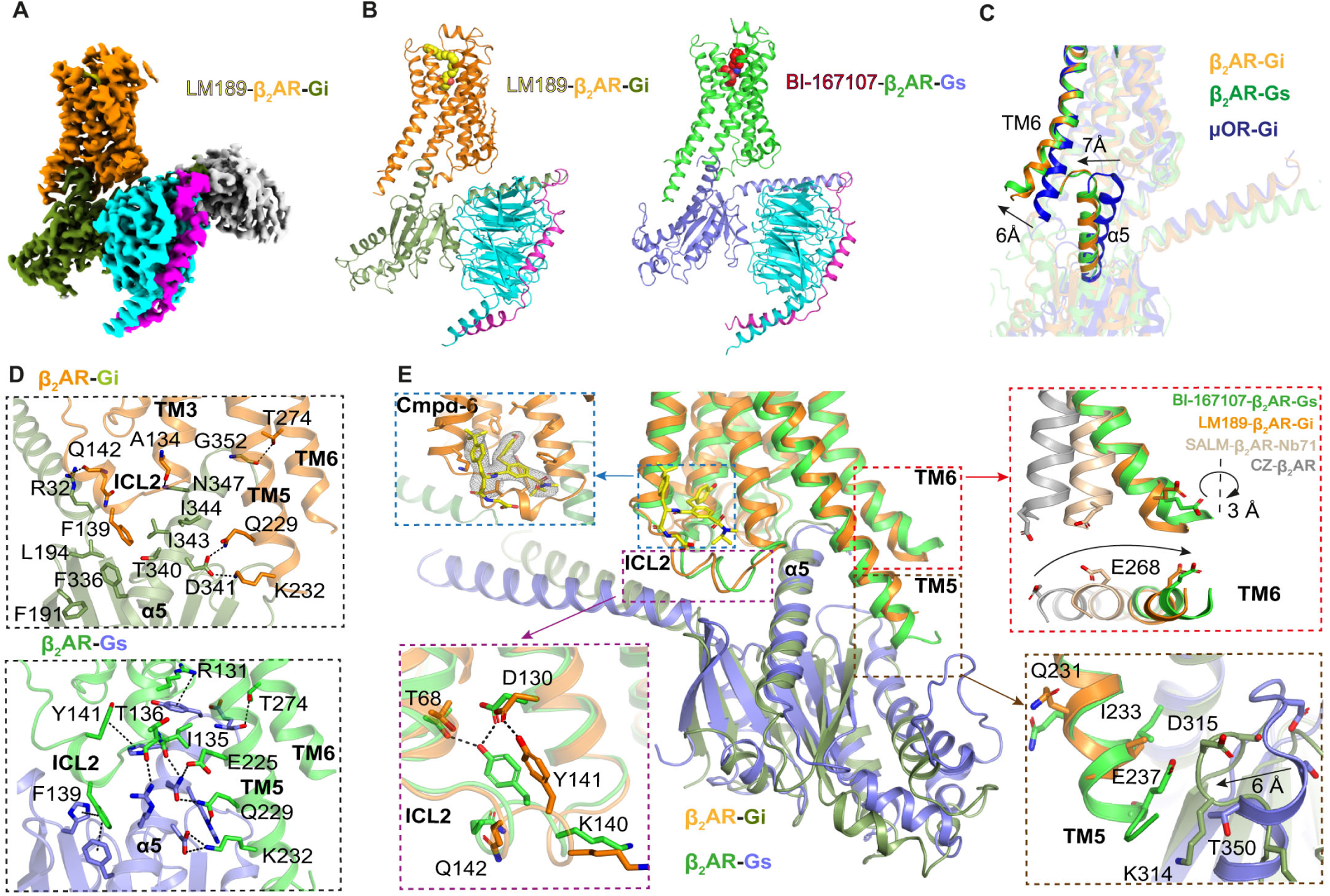
Structure of the LM189-bound *β*_2_AR-Gi complex. **(A)** Cryo-EM density map of the LM189-bound β_2_AR-Gi-scfv16 complex. **(B)** Cryo-EM structure of LM189-bound β_2_AR-Gi complex and crystal structure of BI-167107-bound β_2_AR-Gs (*4*) (3SN6). β_2_AR bound to Gi is colored in orange, Gαi in olive green, Gβ in cyan, Gγ in magenta, scfv-16 in gray. β_2_AR bound to Gs is colored in green, Gαs in slate. **(C)** Relative orientation of receptor TM6 and alpha5 helix of the β_2_AR-Gi (orange), β_2_AR-Gs (green) and µOR-Gi (blue) (*3*) complex structures. **(D)** Top panel: alpha5 engagement of Gi and the interactions formed with the ICL2, TM3 and TM5 of β_2_AR. β_2_AR is colored orange, Gαi in olive green. Lower panel: alpha5 engagement of Gs and the interactions formed with the ICL2, TM3 and TM5 of β_2_AR. β_2_AR is colored in green, Gαs in slate. Polar contacts within 4 Å are highlighted. **(E)** Structural differences between the LM189-bound β_2_AR-Gi complex and BI-167107-bound β_2_AR-Gs (*4*). Top-left blue panel: the PAM cmpd-6FA (*42*), (*43*) (yellow), binding at the top of ICL2. Lower-left magenta panel: when bound to Gi, ICL2 of β_2_AR adopts a slightly different conformation compared to Gs. Top-right red panel: By monitoring TM6 rotation at Glu 268, the TM6 helix is slightly less rotated in the β_2_AR-Gi structure compared to β_2_AR-Gs (∼ 3 Å). Carazolol-bound inactive state β_2_AR is colored in gray (2RH1) (*44*), salmeterol-bound β_2_AR in complex with Nb71 is colored in wheat (6MXT) (*38*), LM189-bound β_2_AR-Gi complex is colored in orange and BI-167107-bound β_2_AR-Gs (3SN6) (*4*) in green. Lower-right brown panel: TM5 of the β_2_AR-Gi complex is one helix turn shorter than TM5 of the β_2_AR-Gs structure, avoiding a steric clash between the tip of TM5 and the connecting loop between the a4 helix and b6 strand of Gi.

Although overall very similar, the β_2_AR-Gi structure is most divergent from the β_2_AR-Gs complex at the intracellular surface, in particular at ICL2, TM6 and TM5 (**Fig. 2E, S3A-F**). To increase complexing efficiency, the β_2_AR-Gi complex was obtained with the aid of the PAM cmpd-6FA, which helps to stabilize ICL2 in the alpha-helical active state conformation (*42*, *43*) (**Fig. 2E, S3B, S3D**). Therefore, our ICL2 structure may differ from that in a complex formed in the absence of cmpd-6FA (discussed further below). We do observe slight differences in the ICL2 active state conformation of β_2_AR-Gi compared to β_2_AR-Gs, in particular at residues Lys 140, Tyr 141, Gln 142 (**Fig. 2E, S3B, S3D**). TM6 of β_2_AR coupled to Gi is very similar to the β_2_AR-Gs structure, with an almost identical outward movement (**Fig. 2E, S3E**). However, by monitoring TM6 helix rotation at residue Glu 268, we observed a slightly smaller outward rotation in the β_2_AR-Gi complex compared to Gs (**Fig. 2E**), resulting in a ∼ 3 Å difference in the outward tilt of Glu 268.

In the β_2_AR-Gs structure, the cytoplasmic end of TM5 undergoes a two-turn elongation compared to the inactive state. In contrast, the intracellular end of the TM5 helix of β_2_AR-Gi elongates only one turn (*4*, *44*) (**Fig. 2E, S3C, S3F**); the cryo-EM density map suggests that this region is highly flexible. This can be attributed to the weaker network of contacts established between TM5 and Gi, mostly consisting of the hydrogen bonds between Asp 341(Gi), Gln 229 and Lys 232 (β_2_AR) (**Fig. 2D**). These interactions are present in the β_2_AR-Gs structure as well, together with three additional hydrogen bonds that help to stabilize the complex (*4*). The shorter TM5 helix prevents a steric clash with the connector loop of the alpha4-beta6 strand of Gi, which moves 6 Å closer to the receptor compared to the corresponding loop in Gs (*3*) (**Fig. 2E, S3F**). Thus, the different TM5 conformations might be important determinants of G protein selectivity and might play a role in the initial steps of G protein coupling.

### LM189 Restricts the Conformational Heterogeneity of the Ligand Binding Pocket

LM189 shares the same tail region as salmeterol, making similar hydrophobic and van der Waals interactions within the β_2_AR exosite (*38*). LM189 also shares the same catechol head group of epinephrine, which forms hydrogen bonds with Asp 113^3.32^, Asn 312^7.39^, Ser 203^5.42^, Ser 207^5.46^ and Asn 293^6.55^ (**Fig. 3A-C**). This polar network has been shown to be important for ligand efficacy at the β_2_AR, and mutations of these residues have an impact on G protein and beta-arrestin recruitment (*38*). We conducted molecular dynamic (MD) simulations to investigate the role of these polar interactions in ligand efficacy and bias. Rotameric analysis of Ser 207^5.46^ and Asn 293^6.55^ showed that ligand-receptor interactions in the orthosteric pocket of LM189-bound β_2_AR were less heterogenous and more stable compared to epinephrine-bound β_2_AR (**Fig. 3D**). Ser 207^5.46^ and Asn 293^6.55^ clearly adopted a favored conformation for LM189-coupled receptor, whereas a broader ensemble of rotamers was sampled in the presence of epinephrine (**Fig. 3D**). For LM189, a very stable hydrogen bond network was formed between the ligand and Ser 203^5.42^, Ser 207^5.46^ and Asn 293^6.55^ (**Fig. 3E**). Moreover, Asn293^6.55^ also formed a stable network with Tyr 308^7.35^ and Ser 204^5.43^ (**Fig. 3E**). In contrast, epinephrine stabilized a weaker polar network, resulting in the loss of the Asn 293^6.55^ and Ser 203^5.42^ interaction with the meta-OH group during our simulations (**Fig. 3E**).

**Fig. 3.**
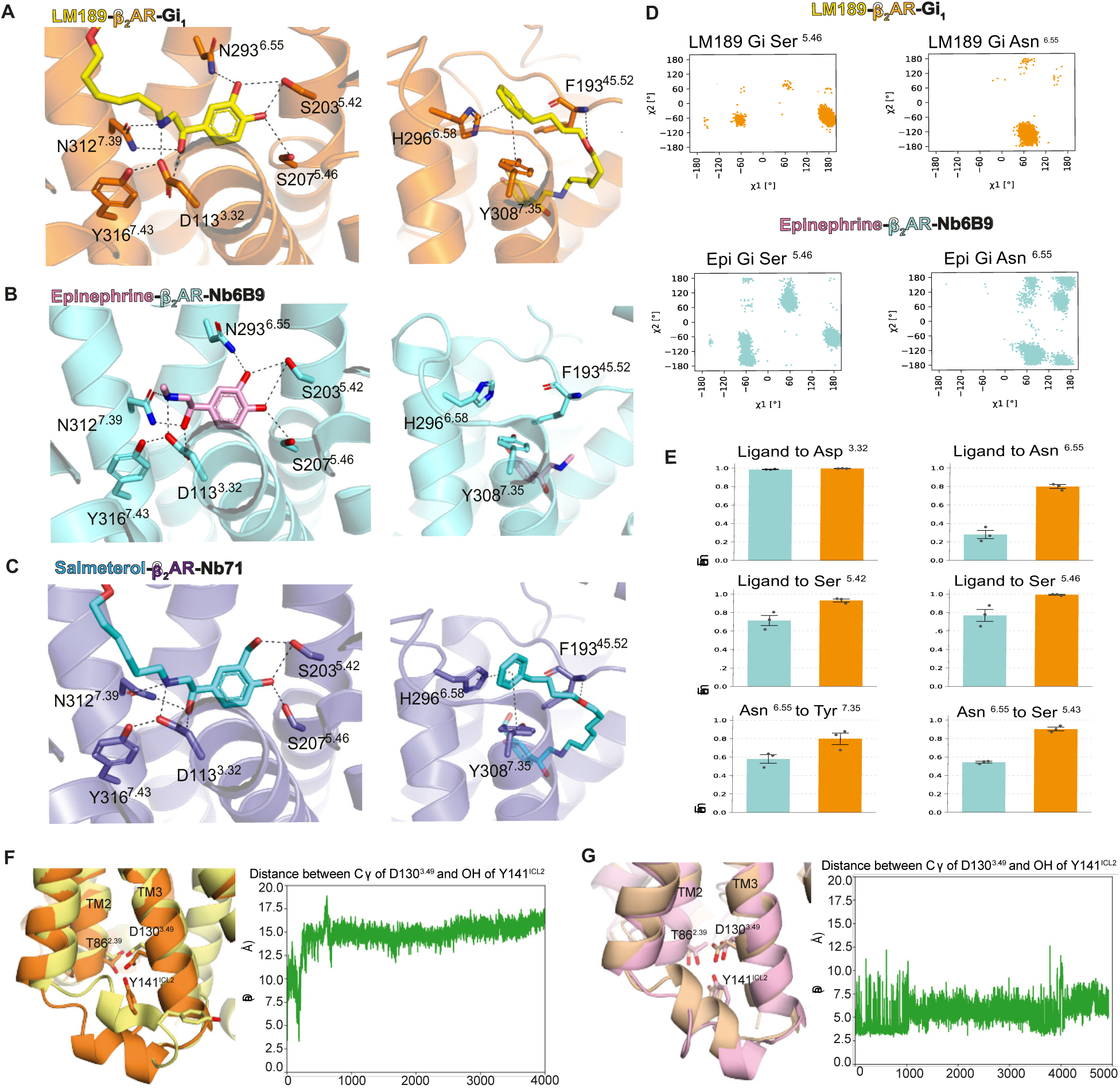
Ligand-binding pocket of the LM189-bound *β*_2_AR-Gi complex. **(A-C)** The orthosteric pocket (left) and exosite (right) of β_2_AR bound to LM189 **(A),** epinephrine **(B)** and salmeterol **(C). (A)** LM189 is colored in yellow, β_2_AR in the β_2_AR-Gi structure is colored in orange. **(B)** Epinephrine is colored in pink, epinephrine-bound β_2_AR in complex with Nb80 (4LDO) (*45*) in aquamarine. **(C)** Salmeterol is colored in blue, salmeterol-bound β_2_AR in complex with Nb71 (6MXT) (*38*) in slate. H-bonds are show as dashed lines. **(D-E)** Molecular dynamic simulations of active state β_2_AR-Gi bound to LM189 or epinephrine. **(D)** Rotamer analysis of Ser^5.46^ and Asn^6.55^ of LM189-bound (orange, top panel) and epinephrine-bound (aquamarine, lower panel) β_2_AR-Gi. **(E)** Histograms represent hydrogen-bond formation frequencies as a fraction of time in three 2µs MD simulations. β_2_AR-Gi is represented in orange, epinephrine-bound β_2_AR-Gi in aquamarine. (**F-G**) MD simulations at the intracellular cavity of β_2_AR. **(F)** Left, comparison of the cryoEM structure of the LM189-bound β_2_AR (orange) and a representative MD snapshot of the LM189-bound β_2_AR (yellow). Right: plot shows the progression of the distance between the Cγ of Asp130^3.49^ and OH of Tyr141^ICL2^ over the course of 4 µs. **(G)** Left, comparison of the epinephrine-bound β_2_AR model (pink, based on the BI-167107-bound β_2_AR crystal structure, PDB: 3SN6) and a representative MD snapshot of the epi-bound β_2_AR model (wheat). Right: plot shows the progression of the distance between the Cγ of Asp130^3.49^ and OH of Tyr141^ICL2^ over the course of 4.9 µs.

Additional MD simulations at the extracellular region of β_2_AR bound to the ligands epinephrine, LM189 and salmeterol indicated a more flexible ECL3 in the epinephrine-bound state coupled to Gs (red box in **Fig. S3G**) compared to LM189-coupled receptor, which displayed similar ECL3 motions to salmeterol-bound β_2_AR (**Fig. S3G**). Additionally, we observed higher flexibility of ECL1 and ECL2 when β_2_AR was coupled to Gi (**Fig. S3G**). Collectively, our MD simulations suggest that LM189 restricts the conformational heterogeneity of the ligand binding-pocket compared to the more flexible configuration in the presence of epinephrine.

Since the β_2_AR-Gi and β_2_AR-Gs structures mostly diverge at ICL2, we conducted MD simulations at the intracellular cavity of the receptor to investigate ICL2 conformational changes in Gs and Gi bound receptors. Simulations of the LM189-β_2_AR-Gi and epinephrine-β_2_AR-Gs complexes after removal of the respective G proteins showed partial unwinding of the ICL2 helical structure (**Fig. 3F, G**). We observed greater ICL2 mobility when the receptor was bound to LM189, as shown by the repositioning of residue Tyr141^ICL2^ (**Fig. 3F, G**). This suggests increased ICL2 dynamics in the absence of cmpd-6FA and G-proteins. Because MD simulations timescales are too short (nsec to µsec) to fully characterize the conformational changes occurring at the intracellular cavity (msec), we decided to conduct additional biophysical measurements to better describe the intracellular rearrangements implicated in G-protein specificity.

### The Role of ICL2 in G protein Specificity

*Hydrogen Deuterium* Exchange Mass Spectrometry (HDX-MS) studies indicate that ICL2 interaction with the hydrophobic core of the G protein alpha subunit is one of the first steps in the receptor-G protein interaction (*13*, *31*, *33*). Previous biophysical and structural investigations have shown that ICL2 of β_2_AR forms a loop in the inactive state (*44*) and a helix in the active-state conformation (*4*). In our β_2_AR-Gi complex, ICL2 forms a partial helix (**Fig. 2D, E**). However, as noted above, this might be due to the presence of the cmpd-6FA PAM, which binds on top of ICL2 and stabilizes the helical conformation (*42*), (*43*). Previous NMR studies suggest that ICL2 does not form a helix when coupled to Gi (*14*). We used continuous wave-electron paramagnetic resonance (CW-EPR) spectroscopy to investigate ICL2 conformational dynamics, to better understand its implication for G protein selectivity at the β_2_AR. The EPR spectral lineshape is sensitive to protein motion on the nanosecond timescale. For properly placed labels, conformational exchange that takes place on the microsecond or longer timescale results in a composite lineshape comprised of the weighted sum of spectral components arising from different conformational states (*46*). We monitored the conformational dynamics at ICL2 by site-directed spin labeling of Q142C, a residue located in the middle of ICL2 which served as conformational reporter (**Fig. 4A**).

**Fig. 4.**
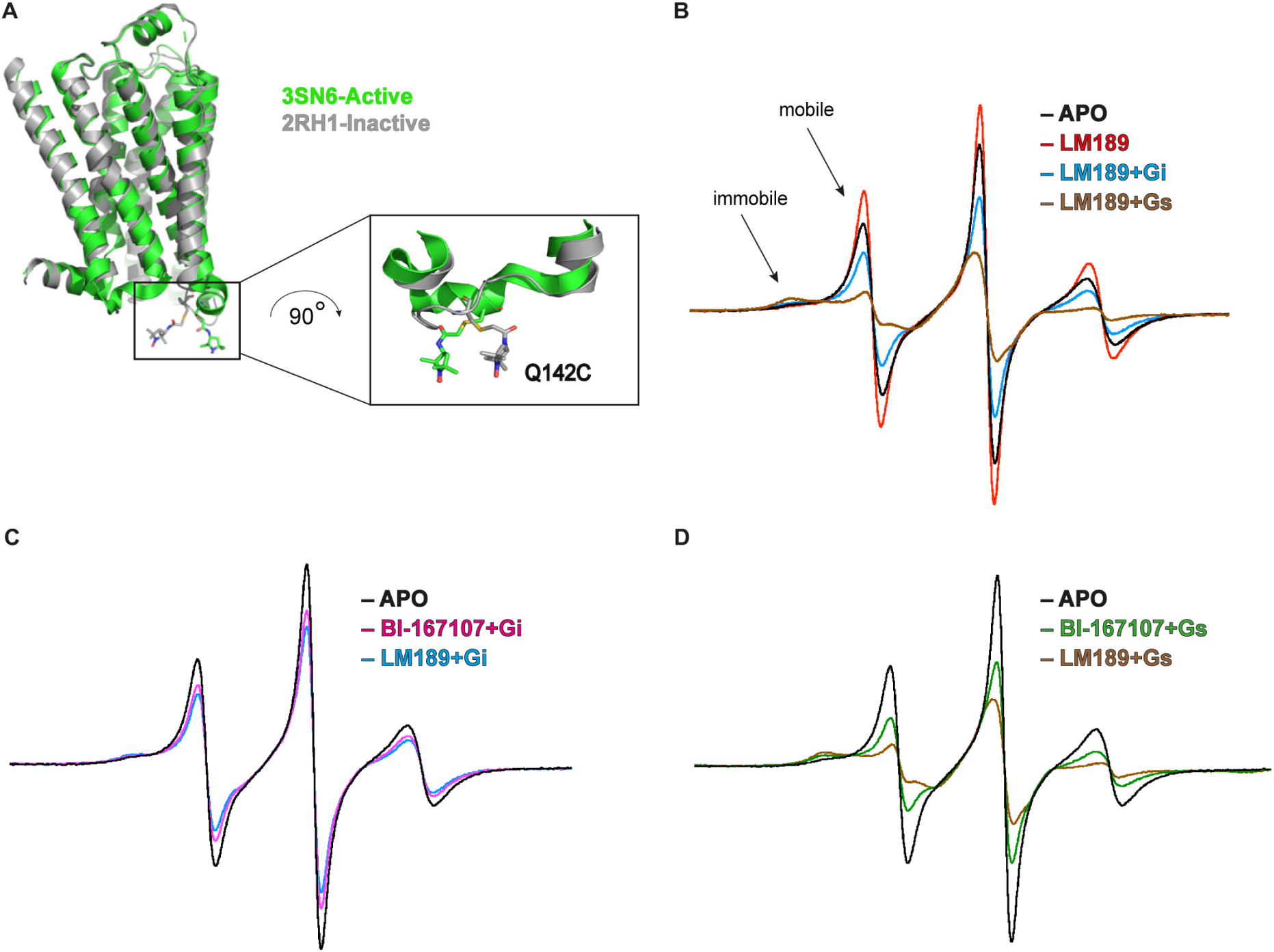
CW-EPR studies of ICL2 of *β*_2_AR. **(A)** A minimal-cysteine version of the β_2_AR with an acetamido-PROXYL spin-label side chain shown at the mutated Q142C residue on ICL2 for EPR studies. Inactive state β_2_AR (2RH1) (*44*) is colored in gray, active state β_2_AR (3SN6) (*4*) is colored in green. **(B)** Superimposed CW-EPR spectra of β_2_AR in the apo, LM189-bound, Gi and Gs-protein bound conditions. The regions of the low-field line dominated by mobile and immobile components are indicated. **(C)** Superimposed CW-EPR spectra in the apo, LM189-bound and BI-167107-bound receptor in complex with Gi. **(D)** Superimposed CW-EPR spectra in the apo, LM189-bound and BI-167107-bound receptor in complex with Gs. All spectra are area-normalized and color-coded as indicated.

We collected CW spectra for the unliganded (apo) receptor and for β_2_AR bound to the biased ligand LM189, the full agonist BI-167107 and the partial agonist salmeterol (**Fig. S4A**). The spectrum of the apo receptor is dominated by a component reflecting the high mobility of the spin label, with a minor component reflecting an immobilized state (**Fig. 4B, S4A**). Based on prior structural evidence, the mobile and immobile components are taken to reflect the loop and helical states of ICL2, respectively. Thus, the CW spectrum of the apo receptor indicates the presence of an equilibrium between a loop and helical state of ICL2 (**Fig. 4B, S4A**). Ligand binding did not substantially change the CW spectral lineshape compared to the apo receptor, except for the case of LM189, where we observed an increase in the population of the mobile spectral component (**Fig. 4B, S4A-B**), in agreement with our MD simulations (**Fig. 3F**). Upon G protein coupling we observed a decrease in the mobile component population and a concomitant increase in the immobile component (**Fig. 4C-D, S4A-D**), a change consistent with ICL2 transition from loop to helix. This shift was greater for Gs than Gi coupling (**Fig. 4C-D**). Interestingly, helix formation upon Gs coupling was more pronounced in the salmeterol and LM189 conditions (**Fig. 4D, S4A, B, D**). This suggests similar conformational dynamics at ICL2 of the receptor bound to LM189 and salmeterol upon Gs coupling.

Taken together, these observations suggest that ICL2 of β_2_AR adopts a stable helix conformation when bound to Gs, while being more dynamic when bound to ligands and Gi. F139 of ICL2 of β_2_AR inserts within the G protein hydrophobic pocket formed by the αN/β1 hinge, β2/β3 loop, and α5 (**Fig. 2D**). The persistence of the unstructured ICL2 conformation upon Gi coupling, when compared with Gs coupling, correlates with the less stable interactions that we observe in the β_2_AR-Gi structure (**Fig. 2D**). Since ICL2 participates in the initial G protein recognition and engagement with the receptor, together with the alpha5 and distal C-terminus of the G protein, different ICL2 conformations might also be associated with structurally distinct mechanisms for primary vs secondary G protein coupling. This agrees with previous NMR and mass spectrometry findings that propose a different role for ICL2 in G protein recruitment for Gs and Gi-coupled receptors (*13*, *14*, *31*, *33*).

### LM189 Stabilizes a Gi-Specific TM6 Conformation

TM6 outward movement has been investigated with a variety of biophysical methods to monitor GPCR activation (*17*, *18*, *32*). TM6 opening in the β_2_AR-Gi complex is very similar to the one observed in the β_2_AR-Gs structure (**Fig. 2E**) and does not explain the increased efficacy of LM189 for Gαi recruitment. We used fluorescence spectroscopy to investigate the conformational dynamics induced by LM189 and other β_2_AR ligands. For these studies, the receptor was labeled with monobromo-bimane at the cytoplasmic end of TM6 (mBBr-β_2_AR) (*47*, *48*) (**Fig. 5A**). As previously observed, salmeterol-bound receptor induced an intermediate TM6 opening, while BI-167107 promoted a larger TM6 movement, which translated into a shift of the probe to a more polar and solvent-exposed environment, characterized by a drop in fluorescence intensity and a red-shift in λ_max_ (*38*) (**Fig. 5B**). Interestingly, LM189-bound receptor showed an even larger decrease in mBBr intensity, suggesting a larger TM6 opening compared to the full agonist BI-167107, or a larger fraction of the receptor in an active conformation (**Fig. 5B**). We observed similar results in experiments performed in lipid nanodiscs (HDL particles, **Fig. S5A**), with a slightly less-pronounced shift in λ_max_ for LM189 compared to BI-167107. This suggests that the TM6 conformation stabilized by LM189 is different in its degree of opening and/or rotation from the conformations previously observed with other β_2_AR agonists.

**Fig. 5.**
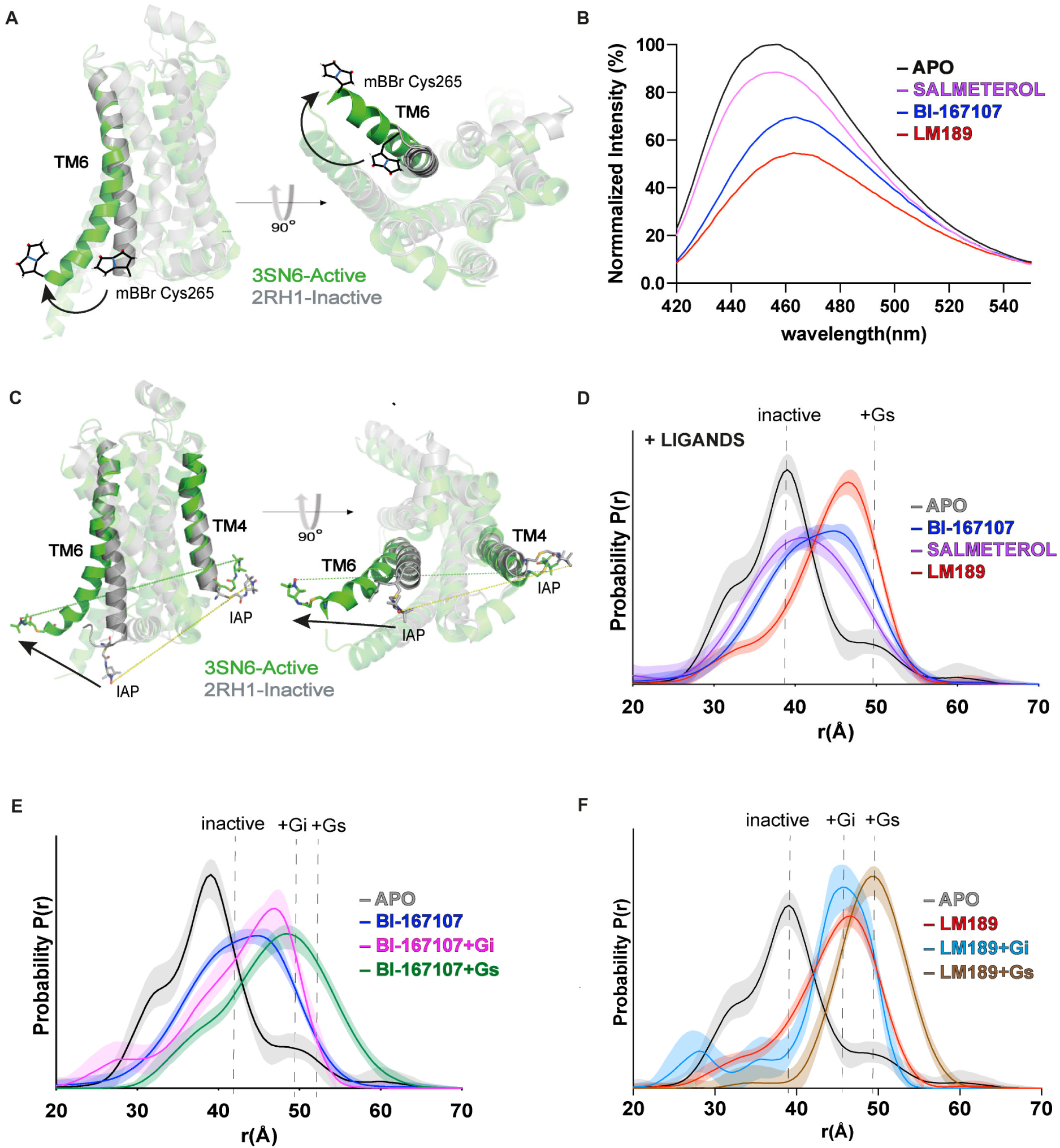
Investigations of TM6 conformational dynamics. **(A-B)** Fluorescence spectroscopy measurements of β_2_AR TM6 conformations. **(A)** Side and intracellular views of β_2_AR labeled with mBBr on Cys265 of TM6. Inactive state β_2_AR (2RH1) (*44*) is colored in gray, active state β_2_AR (3SN6) (*4*) is colored in green. **(B)** Steady-state fluorescence emission spectra of mBBr-labeled β_2_AR purified in LMNG/CHS in the presence and absence of ligands. The spectra are normalized relative to apo (unliganded) receptor (gray). **(C-F)** TM4/6 DEER measurements of β_2_AR purified in LMNG/CHS in the presence of ligands and G proteins. **(C)** Side and intracellular views of the receptor labeling sites on TM4 and TM6. Inactive state β_2_AR (2RH1) (*44*) is colored in gray, active state β_2_AR (3SN6) (*4*) is colored in green. **(D)** Ligand dependence of TM4/6 distance distributions. **(E)** G protein dependence of distance distributions of BI-167107-bound receptor. **(F)** G protein dependence of distance distributions of LM189-bound receptor. Distance distributions are color-coded as indicated.

To better characterize LM189 conformational changes and their role in biased signaling and G protein specificity, we combined double electron-electron resonance (DEER) spectroscopy and single-molecule FRET (smFRET) investigations. For DEER studies, we used a minimal-cysteine version of β_2_AR, spin-labeled with iodoacetoamido-PROXYL (IAP) at the intracellular TM4/TM6 helices (N148C/L266C) (*17*) (**Fig. 5C**). The distance distribution for the unliganded (apo) receptor (gray in **Fig. 5D, S5B-E**) displays two main peaks centered at approximatively 32 Å and 39 Å as well as a smaller peak at 50 Å corresponding to a minor active state population. Addition of the partial agonist salmeterol (purple in **Fig. 5D, S5B, S5E**) broadens the distribution and shifts the most probable distance to ∼41 Å, indicative of a conformationally heterogenous position of TM6 at an intermediate opening relative to both inactive and active distances. As previously reported (*17*), the ultra-high affinity full agonist BI-167107 (blue in **Fig. 5D**, **S5E**) promoted a more open TM6 conformation (∼ 45 Å) but failed to stabilize a fully outward active-state conformation observed in the β_2_AR-Gs structure. Interestingly, consistent with our bimane studies, we observed a greater TM6 outward movement in the presence of LM189, with a distance distribution dominated by a relatively narrow and monomodal peak at ∼ 47 Å, in contrast with the broader distributions measured in the salmeterol and BI-167107 conditions (red in **Fig. 5D, S5E**).

Next, we sought to evaluate the effect of the G proteins Gs and Gi on β_2_AR distance distributions. Gi addition to salmeterol-coupled receptor resulted in a broad distance distribution, only marginally stabilizing the Gi-occupied conformation (**Fig. S5B**). The incomplete shift to fully G-protein occupied receptor observed in the presence of salmeterol is likely due to the partial agonist efficacy of the ligand. In the presence of Gs, salmeterol-bound receptor exhibited a longer most probable distance, ∼ 50 Å, in agreement with the BI-167107-β_2_AR-Gs data (**Fig. S5B, D**). G protein coupling to BI-167107-bound receptor resulted in the stabilization of two distinct distance distributions for Gi and Gs, with most probable distances of ∼ 47 Å and ∼ 48 Å respectively (**Fig. 5E, S5C-E**). These distributions are broader, more multimodal, and present greater probability density in the 30-40 Å range than those for LM189-coupled receptor bound to G proteins. Interestingly, addition of Gi to LM189-coupled β_2_AR (∼ 46 Å most probable distance) (light blue in **Fig. 5F**) populated a distance distribution with a great degree of overlap with that of the LM189-coupled receptor alone (red in **Fig. 5D, 5F, S5E**). In contrast, Gs coupling to LM189-bound β_2_AR resulted in a longer most probable distance (∼ 49 Å) (brown in **Fig. 5F**), corresponding to a further ∼3 Å shift compared to Gi-bound receptor. Based on these observations, LM189 may be Gi-biased because it stabilizes a TM6 outward conformation corresponding to the Gi-competent state, facilitating Gi recruitment. Conversely, Gs stabilizes a longer most probable distance, suggesting that TM6 populates slightly different conformations when the receptor is bound to Gi or Gs (**Fig. 5D-F, S5B-E**).

For smFRET studies, the minimal cysteine β_2_AR construct was labeled at the intracellular TM4/TM6 with the donor and acceptor fluorophores DY549P1 and Alexa647. Labeled β_2_AR was subsequently surface-immobilized and imaged using an objective-TIRF microscope (*49*, *50*). Similar to our DEER results, the apo (unliganded) receptor mainly populated two high-FRET states centered at ∼ 0.9 and 0.7 FRET, corresponding to the close-proximity of helices TM4 and TM6, typical of the inactive receptor (black in **Fig. 6A**, **B**). Individual FRET traces also showed rare excursions to lower FRET states, ranging from ∼ 0.2 to ∼ 0.6 FRET (black in **Fig. 6B**). Also, in agreement with our DEER data, upon addition of the full agonist BI-167107 (blue in **Fig. 6A**, **B**) an intermediate state at ∼ 0.6 FRET became the predominant population, at the expense of the inactive higher FRET states. We also observed a modest increase in the low-FRET values at ∼ 0.2, suggesting the enhancement of the proportion of the receptor in the active state, where the donor and acceptor fluorophores on TM4 and TM6 are further apart (blue in **Fig. 6A**, **B**). The smFRET states observed in the apo and BI-167107 conditions are in agreement with previous smFRET studies (*32*). However, compared to previous smFRET investigations on the β_2_AR (*32*), we were able to resolve transitions between distinct receptor states (inactive, intermediate, active) (**Fig. 6A, B**). This was possible by using the donor/acceptor fluorophores pair DY549P1/Alexa647, which combine a slightly longer Förster radius compared to the previously used Cy3B/Cy7 (*32*) with notably improved signal quality, mostly due to the increased brightness of Alexa647 over Cy7.

**Fig. 6.**
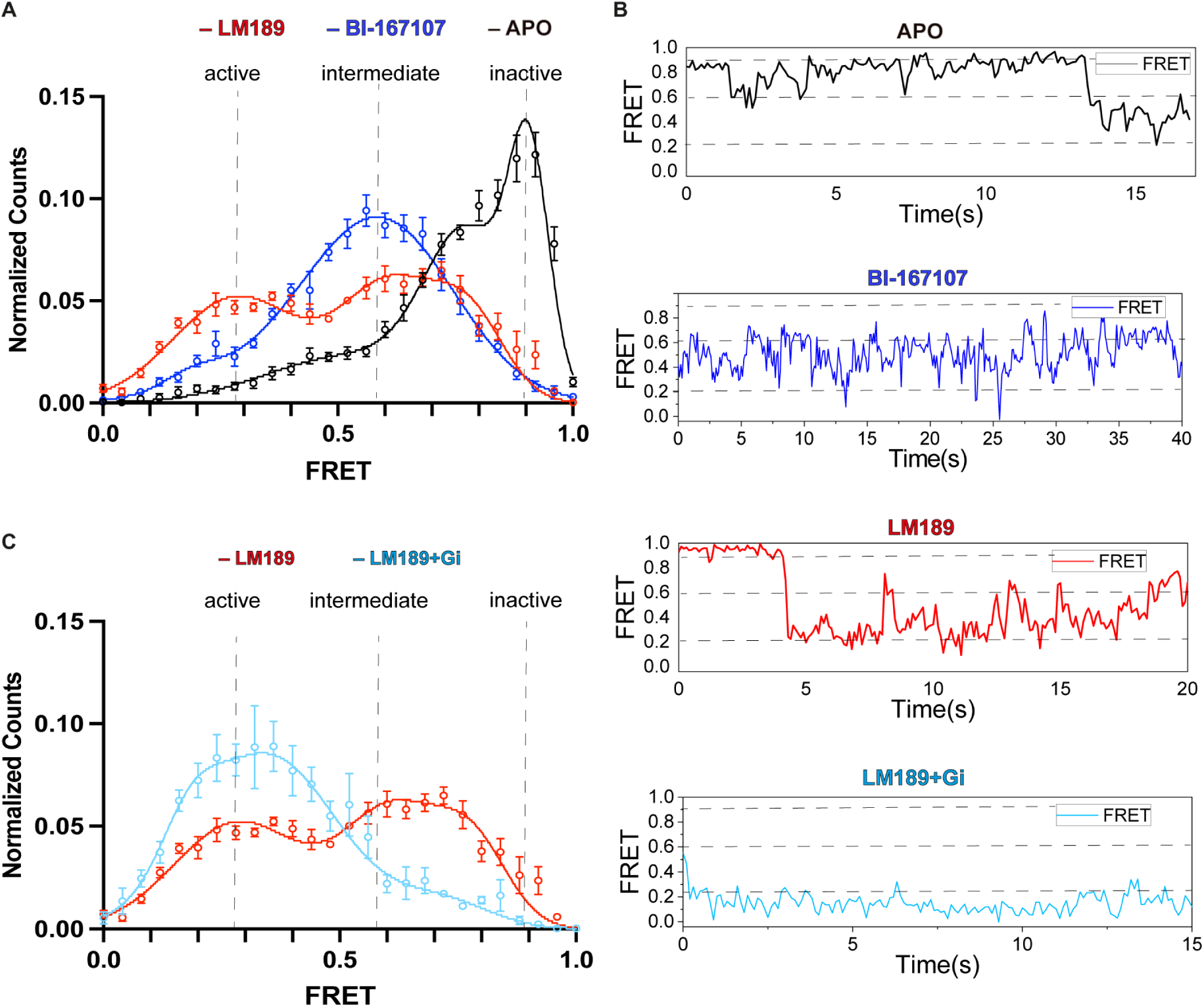
SmFRET distributions of TM6 conformational dynamics. **(A-C)** SmFRET measurements of β_2_AR labeled on TM4/6 with donor and acceptor fluorophores. **(A)** Unliganded (apo, N=269) receptor, in black, mostly populates the inactive states (*∼* 0.9 and 0.7), corresponding to high FRET. BI-167107 (blue, N=265), a β_2_AR full agonist, mostly populates the intermediate state of the receptor (*∼* 0.6) and marginally stabilizes a state at ∼ 0.2 FRET efficiency. LM189-bound receptor (red, N=181) populates the intermediate state at *∼* 0.6 to a smaller degree, in favor of the low-FRET population (∼ 0.3). **(B)** Representative FRET traces and transitions in the apo, BI-167107, LM189 and LM189-bound receptor coupled to Gi and scfv-16. **(C)** Upon Gi coupling, the active state (∼ 0.3), corresponding to low FRET, becomes predominant (light blue, N=105).

Upon addition of the biased agonist LM189, we observed the co-existence of two major FRET populations: the intermediate one at ∼ 0.6 FRET efficiency and a low-FRET state at ∼ 0.3 FRET, which based on our DEER measurements we attribute to the outward TM6 conformation stabilized in the presence of LM189 (red in **Fig. 5D**, **6A**, **B**). The increase in the ∼ 0.3 FRET state was at the expense of the intermediate FRET population, which appeared to be less dominant for LM189-bound receptor compared to the BI-167107 condition (**Fig. 6A**). While BI-167107 only has a minor tail in the low FRET conformation, LM189 populates the ∼ 0.3 FRET state ∼ 40% of the time, even without the addition of G protein (**Fig. 6A**). This conformation corresponds to the Gi-competent state, as observed upon coupling of Gi in complex with scfv-16 to LM189-bound receptor, shifting TM6 to the low-FRET peak (∼ 0.3 FRET) almost completely (light-blue in **Fig. 6B**, **6C**). Collectively, our DEER and smFRET data suggest that LM189 is a Gi-biased ligand because, unlike other agonists, it very effectively stabilizes the Gi-competent conformation, thereby facilitating Gi recruitment. While we observed one main peak in the DEER distance distribution for receptor bound to LM189 (**Fig. 5D**), we detected two populations upon LM189 addition in our smFRET measurements (**Fig. 6A**). It is possible that the two conformations observed by smFRET exhibit subtle differences in TM6 position or local environment and while these motions are amplified by the smFRET environmental sensitive dyes used in this study, they are not observable as TM4-TM6 distance changes in DEER. This is also supported by our ensemble fluorescence experiments (**Fig. 5B, S5A**), indicating that the LM189-stabilized TM6 conformation diverges in its degree of opening and helix rotation from the other ligands.

## Discussion

Despite the large number of structures available of GPCRs in complex with different G protein subtypes (*3–6*), the molecular determinants of G protein coupling specificity remain elusive. This is because specificity is hypothesized to be determined at the level of intermediate receptor conformations, that due to their transient nature are usually not accessible by traditional structural methods. In this context, GPCR agonists that are able to select specific signaling pathways, also called biased agonists, can drive the G protein coupling preference for a specific GPCR.

In this work, we aimed at a better understanding of the role of ligand efficacy and bias for G protein coupling specificity. We studied the binding of β_2_AR to its main G protein, Gαs, and its secondary G protein, Gαi, using a combination of structural and biophysical methods. We identified salmeterol (*38*), a β_2_AR partial agonist for Gs binding, as a full agonist for the recruitment of Gi with an efficacy greater than the native agonist epinephrine (**Fig. 1A-C**). Based on the salmeterol scaffold, we developed LM189, a Gi-biased agonist according to our Glo assay and BRET investigations (**Fig. 1, S1**). For BRET assays, we employed mini-G proteins, engineered G-proteins that only contain the GTPase domains of Gα subunits. While the results obtained by BRET experiments are in agreement with our Glo assay investigations, it is important to highlight that the differences measured with the use of mini-G proteins do not necessarily reflect receptor selectivity in the context of more physiological systems, where fully reconstituted G protein hetero-trimers better reflect receptor specificity profile.

LM189 was used to obtain the cryo-EM structure of the β_2_AR-Gi complex (**Fig. 2, S2**). The orthosteric binding pocket of the LM189-β_2_AR-Gi structure is very similar to the binding pocket of β_2_AR in the presence of salmeterol and epinephrine (**Fig. 3A-C**). This suggests that the G protein specificity mechanism exerted by the biased ligand LM189 is rather mediated by intermediate receptor conformations that involve the core and the intracellular cavity of the receptor. We observed the major differences between the β_2_AR-Gi structure and the previously obtained β_2_AR-Gs complex at the intracellular ICL2, TM6 and TM5 (**Fig. 2E**). Different conformations at ICL2, TM5 and TM6 have been reported in other GPCR-G protein complexes for receptors that can couple to both Gs and Gi, indicating that these intracellular domains might be involved in the initial recognition and engagement of distinct G protein subtypes (*7*, *8*, *33*). In the β_2_AR-Gs complex, we observed a ∼ 3 Å TM6 further opening compared to the Gi-bound structure (**Fig. 2E**). This structural difference has been also measured by DEER (**Fig. 5E, F**). In agreement with previous GPCR-G protein structures, Gs and Gq,11,12,13 coupling is associated with a wider G protein-binding pocket relative to Gi-coupled structures, to accommodate the bulkier C-terminus of the α5 helix of the Gs and Gq isoforms and still allow the interaction with the less bulky Gi α5 helix (*9*, *13*, *51*). Because the size of the G protein binding pocket in the receptor intracellular core may reflect the receptor’s ability to couple to multiple G proteins, it has been hypothesized that receptors that canonically couple to Gs (and Gq,11,12,13) are generally more promiscuous than those that are classified as Gi-coupled(*11*). It should however be noted that the stable nucleotide-free β_2_AR-Gs complex used in cryo-EM studies may contain structural changes in the G protein binding that are not observed in the transient nucleotide-free state in vivo (*52*). In addition, previous smFRET investigations have found evidence for at least one transient intermediate state in the process of complex formation (*32*). Therefore, biophysical studies done in the absence of G-proteins provide more accurate information about the ligand-specific structural changes in the cytoplasmic surface that determine coupling specificity.

To investigate the receptor conformations at the basis of G protein specificity and the role of the biased ligand LM189, we conducted spectroscopic investigations at the intracellular ICL2 and TM6 of the receptor using CW-EPR, fluorescence spectroscopy, DEER and smFRET. To note, these experiments were conducted in the absence of cmpd-6FA, the PAM used for structure stabilization of the β_2_AR-Gi complex. CW-EPR investigations indicate that the loop-helix equilibrium of ICL2 is shifted towards the helical state to a greater degree by Gs binding compared with Gi binding (**Fig. 4B-D, S4B-D**), in agreement with previous NMR investigations (*14*). To note, when β_2_AR was bound to LM189 we observed an increase in the mobile component of the CW-EPR spectra, a conformational transition that might be relevant for the initial stages of G protein recognition (**Fig. 4B, S4B-D**). In agreement with CW-EPR measurements, MD simulations at the LM189-β_2_AR-Gi structure also show ICL2 unwinding upon removal of Gi, indicating that ICL2 conformational transitions may be involved in G-protein coupling specificity (**Fig. 3F, G**).

Fluorescence spectroscopy studies with a probe at the intracellular end of TM6 of β_2_AR suggested the presence of a distinct conformation for LM189-occupied receptor compared to other non-biased agonists, characterized by a larger TM6 outward movement (**Fig. 5B, S5A**). DEER experiments showed that LM189-coupled β_2_AR populates a strikingly similar TM4/TM6 distance distribution to that observed upon addition of the G protein Gi, suggesting that LM189 stabilizes the same receptor TM6 conformation as Gi (**Fig. 5D**, **5F**). This may explain the preferential Gαi recruitment observed in our experimental data by the biased ligand LM189 (**Fig.1, S1**). In agreement with our structural work, which revealed a difference in rotation of the intracellular end of TM6 in the β_2_AR-Gs structure compared to β_2_AR-Gi (**Fig. 2E**), DEER measurements showed an additional ∼ 3 Å TM6 structural change upon Gs binding (**Fig. 5F**), indicating slightly different conformations for receptor bound to different G protein subtypes (**Fig. 5F, S5B-D**). By smFRET, we detected a predominant intermediate state (∼ 0.6 FRET) for the TM4-TM6 labeling sites of β_2_AR in the presence of the full agonist BI-167107 (**Fig. 6A**, **6B**). In contrast, we observed two FRET states for receptor bound to LM189, the intermediate FRET state (∼ 0.6 FRET) and a low-FRET state (∼ 0.3 FRET), the latter populated even in the absence of the G protein (**Fig.6A**, **6B**). Binding of the G protein Gi shifted the equilibrium of LM189-bound receptor towards the ∼ 0.3 FRET population (**Fig. 6B**, **6C**), corroborating our finding that LM189 is a Gi-biased agonist for its ability to stabilize the Gi-competent conformation.

In this study, we observed that, relative to non-biased agonists, the biased ligand LM189 stabilizes a distinct, specific TM6 conformation (**Fig. 5B, 5D**, **6A**) and increases ICL2 dynamics (**Fig. 4B-D**). Altogether, our investigations reveal that distinct conformations at ICL2 and TM6 of β_2_AR are required for the binding of the different G protein subtypes Gαs and Gαi, underlying the importance of receptor conformational dynamics for coupling specificity and promiscuity. Altogether, these results deepen our understanding of G protein specificity and bias and can be useful in the design of ligands that select for preferred signaling pathways.

## Materials and Methods

### Expression and purification of the *β*_2_-adrenergic receptor (*β*_2_AR)

The β_2_AR construct named PN1 was used for all experiments except EPR, DEER and smFRET investigations. PN1 was expressed and purified as previously described(*35*). Briefly, receptor was expressed in *Spodoptera frugiperda (Sf9)* insect cells (Expression Systems, cell line IPLB-Sf-21-AE, catalog number 94-001S) using the baculovirus method, and media was supplemented with 1 μM alprenolol. Cells expressing β_2_AR were harvested and lysed as previously described(*35*). The receptor was solubilized from membranes using 20 mM hydroxy-ethylpiperazine ethane sulfonic acid (HEPES), pH 7.4, 100 mM sodium chloride (NaCl), 1% n-dodecyl-β-D-maltopyranoside (DDM), 0.03% cholesteryl hemisuccinate (CHS), 2 mM MgCl_2_, 1 μM alprenolol and protease inhibitors. Membranes were homogenized with a douncer and the soluble fraction was isolated by centrifugation and applied to a M1 anti-FLAG immunoaffinity resin. The receptor bound to the resin was extensively washed with 20 mM HEPES pH 7.4, 350 mM NaCl, 0.1% DDM, 0.01% CHS, 10 μM alprenolol, protease inhibitors to lower the detergent concentration. To exchange detergent from 0.1% DDM/0.01% CHS to 0,01% (w/v) lauryl maltose neopentyl glycol (LMNG, Anatrace)/0.001% CHS, the receptor was extensively washed with a progressive gradient of DDM: LMNG buffer. In parallel, while the receptor was bound to the resin, alprenolol was removed by washing with saturating concentrations of the low affinity antagonist atenolol. Due to the fast dissociation kinetics of atenolol from the β_2_AR, subsequent washes with ligand-free buffer yielded unliganded β_2_AR. The receptor was then eluted in a buffer consisting of 20 mM HEPES pH 7.4, 150 mM NaCl, 0.01% LMNG/0.001% CHS, FLAG peptide and 5 mM EDTA. Receptor was further purified by size exclusion chromatography (Superdex 200 10/300 gel filtration column) in buffer containing 20 mM HEPES pH 7.4, 150 mM NaCl, 0.01% LMNG/0.001% CHS. Finally, β_2_AR was concentrated to 250uM, flash frozen after addition of 20% glycerol and stored in -80°C. As we did not perform an additional purification step using alprenolol resin, the functional fraction of purified receptor was assessed by direct coupling of purified β_2_AR to the cognate G protein Gs. Immediately after purification and size exclusion chromatography, PN1 was labeled with *monobromo*(trimethylammonio)*bimane (mBBr) in the presence of* 100 μM tris(2 - carboxyethyl)phosphine (TCEP). Excess dye was removed by size exclusion chromatography and the labeled receptor was incubated with 10-X molar ratio of the agonist ligand *BI-*167107 for 30 minutes. Subsequently, a 1:1 ratio of Gs protein was added for 1 hour, followed by overnight treatment with apyrase (1-unit, NEB) on ice. Following size exclusion chromatography, 80% of the receptor purified in 0.1% DDM, 0.01% CHS and 90% of β_2_AR in 0.01% LMNG/0.001% CHS was attested to be functional, as capable to bind to Gs and to run as a single mono-disperse peak on SEC.

To favor Gi coupling, we avoided working with buffers containing CHS(*35*). To perform the experiments in this work, β_2_AR was extensively diluted into buffers not containing CHS or reconstituted onto HDLs along with neutrally charged lipids. For Cryo-EM purposes, β_2_AR was purified in 20 mM HEPES pH 7.4, 150 mM NaCl, 0.1% DDM, 0.01% CHS as previously described and subsequently exchanged to LMNG/cholesterol micelles to avoid the use of CHS. 5% (w/v) LMNG and 2mol% cholesterol was prepared by OVN stirring followed by sonication in buffer containing 200mM HEPES, pH 7.4. While bound to M1-Flag during purification, β_2_AR was buffer exchanged onto 20 mM HEPES pH 7.4, 150 mM NaCl, 0.01%LMNG/0.003%cholesterol micelles. After the exchange, the receptor was washed with atenolol and eluted from the M1-Flag. All subsequent steps took place as previously described, using LMNG/cholesterol buffers instead of LMNG/CHS.

### Expression, purification and labeling of the *β*_2_AR for CW-EPR and DEER experiments

For CW-EPR experiments we used the β_2_AR minimal cysteine background construct called β2Δ5, which has the following mutations: C77V, C265A, C327S, C378A, and C406A(*17*). In addition, β2Δ5 has an N-terminal FLAG sequence and a hexa-histidine sequence at the C-terminal, as well as methionine substitutions M96T and M98T to increase expression levels.

For DEER investigations, we used the β_2_AR minimal cysteine background construct β2Δ6, which carries an extra mutation at C341L, removing the palmitoylation site. Cysteine mutations at desired locations were re-introduced in the β2Δ5 and β2Δ6 backgrounds to site-specifically label these constructs for CW-EPR and DEER. To monitor ICL2 rearrangements by CW-EPR, Q142C on ICL2 was inserted into a β2Δ5 background. For DEER investigations, we used the previously established β2Δ6 148C/266C construct, with reintroduced cysteines at residues N148C and L266C(*32*). The single cysteine mutants N148C and L266C were also produced for control experiments to monitor nitroxide probe mobility and optimal labeling conditions.

All β2Δ5 and β2Δ6 constructs used for EPR and DEER were cloned into the pcDNA-Zeo-tetO vector as previously described(*22*). Constructs were transfected into the suspension cell line tetracycline-inducible Expi293^TM^ cells (Thermo, catalog number A14635), stably expressing the tetracycline repressor(*53*). Expifectamine transfection kit (Thermo) was used to transfect the cells according to the manufacturer’s recommendations. 2 days post-transfection, receptor expression was induced with doxycycline (4 mg/mL, 5 mM sodium butyrate) in the presence of 2 µM alprenolol. 30 hours post induction the cells were harvested by centrifugation and the pellet was frozen in liquid nitrogen and stored at -80°C. The receptor was subsequently purified in DDM/CHS and exchanged to LMNG/CHS as described above. After elution from the M1-Flag column, the receptor was labeled with the spin label reagent iodoacetamido-PROXYL (IAP) in the presence of 100 μM tris(2-carboxyethyl)phosphine (TCEP) in buffer containing 20mM Hepes pH 7.4, 150mM NaCl, 0,01%LMNG/0,001% CHS. 25-fold molar excess of IAP was added to 40 µM β2Δ6 receptor for 3 hours at RT. For β2Δ5, 10-fold molar excess of IAP was added for 2 hours at RT. After quenching of the reaction with 5mM final L-cysteine, the receptor was separated from the excess spin label by size exclusion chromatography (Superdex 200 10/300 gel filtration column) in SEC buffer (20 mM HEPES pH 7.4, 150 mM NaCl, 0.01% LMNG) prepared with D_2_O and not containing CHS.

### Expression, purification and labeling of the *β*_2_AR for smFRET experiments

The β2Δ6 148C/266C construct was used for smFRET experiments. Expression and purification of β2Δ6 148C/266C was carried out as described for the DEER experiments. For smFRET labeling, receptor eluted from M1-Flag was diluted to 10uM in 20mM Hepes pH 7.4, 150mM NaCl, 0,01%LMNG/0,001% CHS and 100 μM tris(2 -carboxyethyl)phosphine (TCEP) was added for 20min. Labeling was conducted in the presence of 2uM atenolol and was initiated by addition of 10-X of the pre-mixed fluorescent dye pair DY559P1 and Alexa647, at 1:1 ratio. After one hour at RT, the reaction was quenched with 5mM L-cysteine. Further sample purification, TCEP and dye removal was performed by size exclusion chromatography (Superdex 200 10/300 gel filtration column) in 20 mM HEPES pH 7.4, 150 mM NaCl, 0.01% LMNG buffer. Pulled fractions were concentrated to 15uM and froze in liquid nitrogen with 20% glycerol for subsequent smFRET experiments.

### Expression, purification and labeling of the *β*_2_AR receptor for fluorescence spectroscopy

We used the PN1 construct for steady-state ensemble fluorescence experiments. PN1 was expressed in *sf9* cells, purified and exchanged to LMNG as previously described(*35*). For labeling, 100 μM tris(2 -carboxyethyl)phosphine (TCEP) was added for 20min to 10uM PN1 in 20mM Hepes pH 7.4, 150mM NaCl, 0,01%MNG buffer. Labeling was initiated by addition of 20x *monobromo*(trimethylammonio)*bimane (mBBr) for 45 minutes at RT. The reaction was quenched with excess of L-cysteine and the sample was further purified by* size exclusion chromatography (Superdex 200 10/300 gel filtration column) in 20 mM HEPES pH 7.4, 150 mM NaCl, 0.01% LMNG buffer. *Purified and labeled* β_2_AR fractions were *concentrated and flash-frozen for subsequent experiments or for reconstitution into HDLs*.

### HDLs reconstitution

β_2_AR was reconstituted into high-density lipoprotein particles (HDLs) as previously described(*35*). Briefly, receptor was mixed with the MSP1D1 belt protein in a 1:10 receptor:MSP molar ratio and with lipids (3:2 POPC/POPE or POPC/POPG) in a 1:40 MSP:lipids molar ratio. After 2h incubation at 4°C, Biobeads (Biorad) were added at a ratio of 1:10 lipids:beads and incubated for 4h at 4°C to remove the detergent. Upon Biobeads removal by centrifugation, empty discs were separated from β_2_AR-containing discs by M1-flag affinity chromatography and subsequent size exclusion chromatography (Superdex 200 10/300 gel filtration column) in 20 mM Hepes pH 7.4, 100 mM NaCl. HDLs were concentrated to 30uM, flash-frozen in liquid nitrogen and stored at -80°C for future use.

### Expression and purification of heteromeric Gα_1_β_1_γ_2_

Heterotrimeric Gi was expressed and purified as previously described(*3*) with some modifications. Briefly, heterotrimeric Gi was expressed in *Trichoplusia ni (T.ni)* insect cells using the baculovirus method (Expression Systems, Catalog number 94-002S). Two viruses were used to infect the insect cells, one encoding the wild-type human Gα_1_ subunit and another one encoding the wild-type human β_1_γ_2_ subunits. The cells were harvested 48 hours post-transfection and the pellet was flash frozen with liquid nitrogen and stored at -80°C. Cells lysis was conducted in 10 mM Tris, pH 7.4, 100 mM magnesium chloride (MgCl_2_), 5 mM β-mercaptoethanol (βME), 20 mM GDP and protease inhibitors. Membranes were isolated by centrifugation and solubilized using a douncer in 20 mM Hepes pH 7.4, 100 mM NaCl, 1% sodium cholate, 0.05% DDM, 5 mM magnesium chloride, 5 mM βME, 20 mM GDP, 20 U calf intestinal alkaline phosphatase and protease inhibitors. After addition of 20mM imidazole, the solubilization mixture was stirred for one hour at 4 degrees. After centrifugation, the supernatant was loaded on a Ni-NTA chromatography column, extensively washed in 0,05% DDM buffer in order to remove the cholate. Subsequently, a progressive gradient of DDM: MNG buffer was used to exchange the detergent from DDM to 0,05% LMNG. After elution in the presence of 250mM imidazole, the purified heterotrimer was de-phosphorylated by lambda protein phosphatase (NEB), calf intestinal phosphatase (NEB), and antarctic phosphatase (NEB) in the presence of 1mM MnCl_2_ for 1h on ice. Gi heterotrimer was separated from excess betagamma using a MonoQ 10/100 GL column (GE Healthcare). The protein was diluted to lower the imidazole concentration and loaded onto the column in 20 mM Hepes pH 7.4, 50 mM NaCl, 1mM MgCl_2_, 0.05% LMNG, 100 mM TCEP and 20 mM GDP. The heterotrimer was eluted with a linear gradient of 0–50% with buffer containing 1M NaCl. Eluted fractions were concentrated to 200-250uM and after addition of 20% glycerol the protein was flash frozen and stored at -80°C.

### Development and synthesis of LM189

#### NMR analyses

The analytical characterization of LM-189 was performed by ^1^H-NMR at 600 MHz and ^13^C-NMR at 151 MHz. Determination of chemical shifts (ppm) was done in relation to the solvent used (CD_3_OD). The NMR samples were measured under nitrogen atmosphere to avoid decomposition of the catecholaminergic product. The usual abbreviations are used for signal multiplicities: s (singlet), d (doublet), t (triplet), q (quartet), quint (quintet), sext (sextet), sept (septet). Coupling constants are given in Hertz (Hz).

#### Polarimetry

Specific optical rotations were measured using a JASCO P-2000 polarimeter. Chamber path length: 100 mm, chamber volume: max. 1.2 mL. The target compound was measured in the form of a clear solution in methanol.

#### UHPLC-MS

LCMS analyses were conducted using a Dionex UltiMate 3000 UHPLC system with a RS diode array detector for the wavelengths 220 nm and 254 nm. A binary solvent system (mixture of 0.1% aqueous formic acid and methanol) was used as the eluent. <u>Column:</u> Kinetex C8 (75 x 2.1 mm, 2.6 µm, 0.3 mL/min flowrate) or ZORBAX ECLIPSE XDB-C8 (100 x 3.0 mm, 3.5 µm, 0.4 mL/min flowrate). Mass detection was performed on a Bruker Amazon SL mass spectrometer (electron spray ionization, ESI).

#### High-resolution MS

HRMS measurements were performed on a Bruker timsTOF Pro device.

#### Preparative RP-HPLC

The target compound was purified by preparative, reverse-phase HPLC, applying mixtures of 0.1% aqueous trifluoroacetic acid and acetonitrile as organic component. The separations were conducted on a preparative HPLC AGILENT SERIES 1100 system equipped with a variable wavelength detector (VWD), on an AGILENT HPLC 1260 Infinity system with a VWD or on an AGILENT HPLC 1260 Infinity II system with a VWD. <u>Column</u>: ZORBAX ECLIPSE XDB-C8 (150 x 21.5 mm, 5 µm).

#### Analytical RP-HPLC

Analytical HPLC runs for purity control were conducted on an AGILENT 1200 series with DAD detector and peak detection at 220, 230, 254 and 280 nm. The employed column was a ZORBAX ECLIPSE XDB-C8 (100 x 4.6 mm, 5 µm) with a flowrate of 0.5 mL/min. The column thermostat was set to 20 °C to obtain uniform results. The binary solvent systems either consisted of 0.1% aqueous TFA + acetonitrile or 0.1% aqueous TFA + methanol. The following two eluent systems were used:

<u>System 1A:</u> 0.1% aqueous trifluoroacetic acid + acetonitrile, 0.5 mL/min: 5% acetonitrile from 0 to 3 min, to 95% at 18 min, 95% from 18 to 24 min, to 5% at 27 min, 5% from 27 to 30 min.

<u>System 1B:</u> 0.1% aqueous trifluoroacetic acid + methanol, 0.5 mL/min: 5% methanol from 0 to 3 min, to 95% at 18 min, 95% from 18 to 24 min, to 5% at 27 min, 5% from 27 to 30 min.

### (*R*)-4-(1-hydroxy-2-((6-(4-phenylbutoxy)hexyl)amino)ethyl)benzene-1,2-diol x TFA (LM-189)

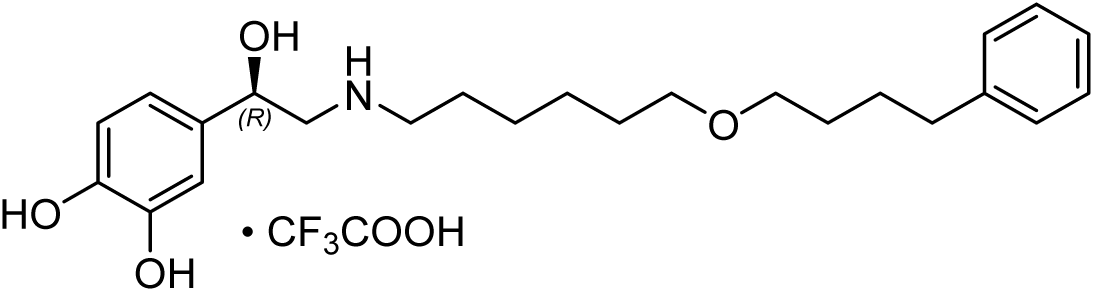

(4-((6-bromohexyl)oxy)butyl)benzene (66.0 mg, 0.21 mmol) was dissolved in DMSO (1 mL) in a microwave tube. To this solution, (*R*)-norepinephrine freebase (100 mg, 0.59 mmol) was added. The vial was set under nitrogen atmosphere, sealed with an aluminum crimp cap and heated to 70 °C for 7 h under light protection. Then, the reaction was stirred at r.t. overnight (18 h) and directly added to a stoichiometric excess of 0.3% TFA solution. The mixture was frozen and lyophilized. The crude compound was purified by preparative HPLC (ZORBAX ECLIPSE XDB-C8, 0.1% TFA + 10% acetonitrile to 95% acetonitrile at 10 min, peak eluted at 7 min) to give the target compound **LM-189** as a yellow-orange oil (59.0 mg, 55% yield).

*Analytical data were in agreement with the literature (racemate of **LM-189**).*(*54*),(*55*)

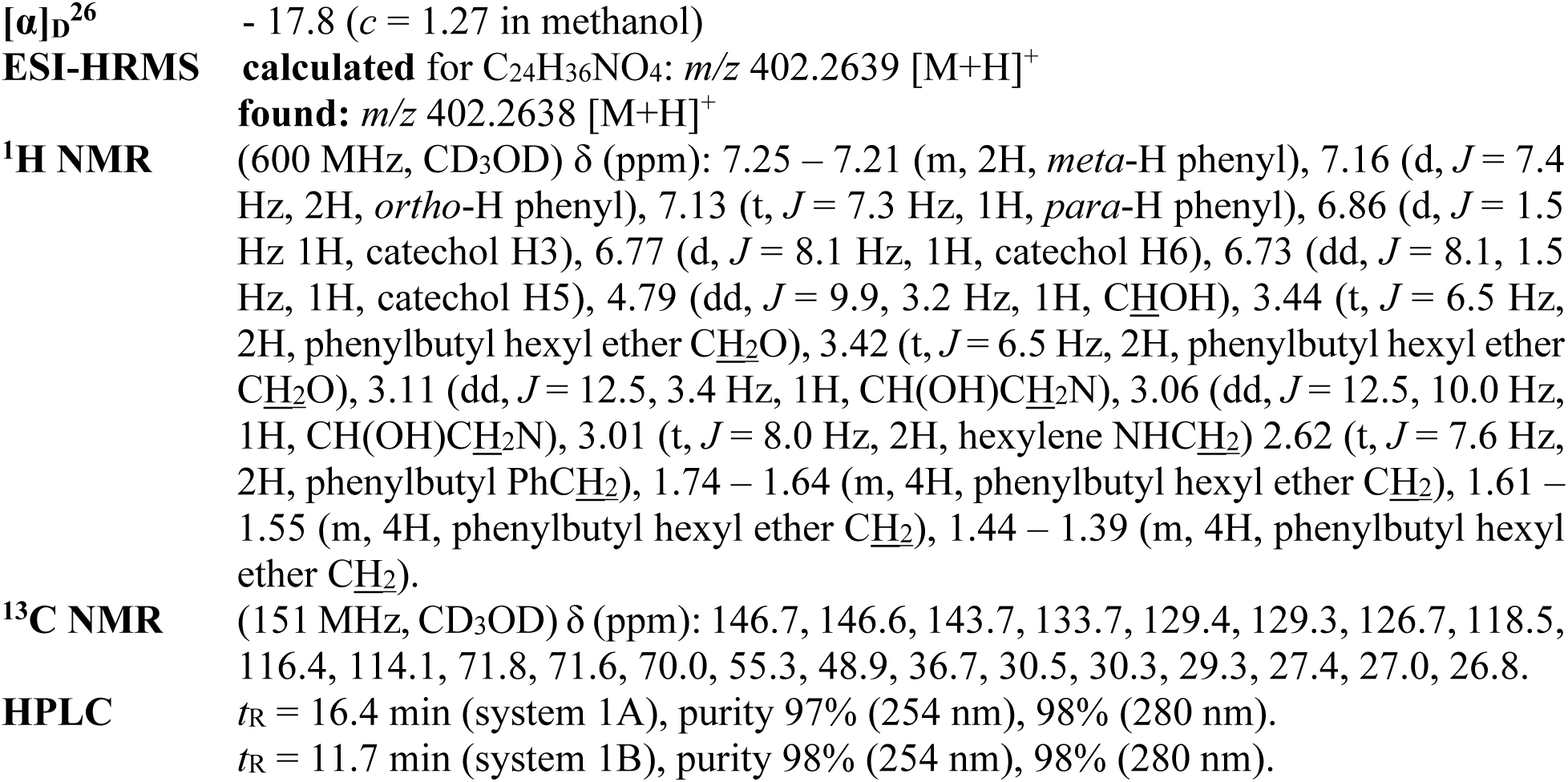

### GTP turnover assay

The GTP turnover assay was adapted from the GTPase-Glo^TM^ assay (Promega) as described previously(*32*). To monitor Gs turnover, PN1 in 0.01%LMNG (75nM final) was incubated for 1 hour at RT with 10-X ligand excess in in 20 mM Hepes pH 7.5, 100 mM NaCl, 0.01% MNG, 20 uM GTP. When complexing with Gi, PN1 was used at 1uM final concentration. For HDLs experiments, we used 300nM receptor in discs in 20 mM Hepes pH 7.5, 100 mM NaCl, 20 uM GTP. To start the reaction, G protein (1uM for Gi, 0.5uM for Gs) in buffer containing 20 mM Hepes pH 7.4, 100 mM NaCl, 20 mM MgCl_2_, 200 uM TCEP, 0.01% LMNG, and 20 uM GDP was added to receptor in MNG. For HDL experiments, we used 0.5uM G protein in 20 mM Hepes pH 7.4, 100 mM NaCl, 20 mM MgCl_2_, 200 uM TCEP, 0.04% DDM, 20 uM GDP. The reaction was carried over for 1 hour to monitor Gi turnover and for 10 minutes for Gs. After incubation at RT, GTPase-Glo reagent supplemented with 10 mM adenosine 5′-diphosphate (ADP) was added to the reaction and incubated for 30 min at RT. Detection reagent was then added and incubated for 10 min at RT prior to luminescence detection using a MicroBeta^2^ microplate counter (PerkinElmer). Experiments were performed as biological triplicates and results were plotted using GrapPad Prism. P values were calculated using the unpaired t test analysis on GraphPad Prism, assuming Gaussian distributions. Ns=(P>0.05), * (P≤0.05), ** (P≤0.01), *** (P≤0.001), **** (P≤0.0001).

### Radioligand binding assay with membranes from HEK cells

Binding affinities towards the human β_1_AR and β_2_AR were determined as described previously(*56*, *57*). HEK293T cells were transiently transfected with the cDNA for β_1_AR and β_2_AR (obtained from the cDNA Resource Center, www.cdna.org). Membranes were prepared showing receptor densities of 3.2 pmol/mg protein (*B_max_* for β _1_AR) and 2.3±0.64 pmol/mg protein (*B_max_* for β _2_AR) and binding affinities for the radioligand [³H]CGP12,177 (specific activity 51 Ci/mmol, PerkinElmer, Rodgau, Germany) of 0.070 nM (*K_D_* for β_1_AR) and 0.095±0.02 nM (*K_D_* for β_2_AR). Competition binding experiments were performed by incubating membranes in binding buffer (25 mM HEPES, 5 mM MgCl_2_, 1 mM EDTA, and 0.006% bovine serum albumin at pH 7.4) at a final protein concentration of 3-10 µg/well, together with the radioligand (final concentration 0.2 nM) and varying concentrations of the competing ligands for 60 minutes at 37 °C. Non-specific binding was determined in the presence of unlabeled CGP12,177 at a final concentration of 10 µM. Protein concentration was established using the method of Lowry(*58*). For data analysis the resulting competition curves were analyzed by nonlinear regression using the algorithms implemented in PRISM 10.0 (GraphPad Software, San Diego, CA) to provide an *IC_50_* value, which was subsequently transformed into a *K_i_* value employing the equation of Cheng and Prusoff(*59*) (Table S1). Mean *K_i_* values were calculated from 7-11 single experiments each performed in triplicates.

### *β*-Arrestin-2 recruitment (PathHunter Assay)

Determination of receptor-stimulated β-arrestin-2 recruitment was performed applying the PathHunter assay (DiscoverX, Birmingham, U.K.), which is based on the measurement of fragment complementation of β-galactosidase as described(*56*, *60*). In detail, HEK293T cells stably expressing the enzyme acceptor (EA) tagged β-arrestin-2 fusion protein were transfected with the cDNA for β_1_AR and β_2_AR, each fused to the ProLink-PK1 fragment for enzyme complementation and transferred into 384 well microplates. Measurement started by incubating cells with epinephrine or LM189 for 90 min and was stopped by addition of the detection mixture. Chemoluminescence was monitored with a Clariostar plate reader (BMG, Ortenberg, Germany) and analyzed by normalizing the raw data relative to basal activity (0%) and the maximum effect of norepinephrine (100%). Dose-response curves were analyzed applying the algorithms for four-parameter non-linear regression implemented in Prism 10.0 (GraphPad LLC, CA) to yield *EC_50_* and *E_max_* values (Table S1). Mean *EC_50_* and *E_max_* values were calculated from 6-17 independent experiments each conducted in duplicates.

### Formation and purification of the *β*_2_AR-Gi-scfv16 complex for Cryo-EM

PN1 in 0,01%LMNG/2mol% cholesterol was incubated with 7-fold molar excess of LM189 ligand for 30 mins at RT. Cmpd-6FA PAM(*42*),(*43*) was then added at 7-fold molar excess and incubated for additional 30 mins at RT. A 1.3-fold molar excess of Gi was added together with 100uM TCEP and incubated for 2h at RT. Afterward, 2-fold molar excess of scfv16 was added to the complex and incubated for 1.5h on ice. Apyrase (1-unit, NEB) was then added, and the complex was incubated OVN on ice. The following day, the complex was diluted in 20mM Hepes pH 7.4, 100mM NaCl, 0,01% MNG/2mol% cholesterol, 0,0033% GDN, 10uM LM189, 1uM cmpd-6FA, 3mM Ca^2+^ and loaded onto M1 anti-FLAG affinity chromatography. Detergent concentration was lowered by washing with buffer containing 20mM Hepes pH 7.4, 100mM NaCl, 0,001% MNG/2mol% cholesterol 0,00025% GDN, 2mM Ca^2+^, 10uM ligand, 1uM cmpd-6FA. Complex was eluted in 20mM Hepes pH 7.4, 100mM NaCl, 0,00075% MNG/2mol% cholesterol, 0,00025% GDN, FLAG peptide, 5mM EDTA, 10uM LM189, 1uM cmpd-6FA. 100uM TCEP, 3mM MgCl2, 1:1 (Gi:scfv) molar ratio of scfv16 were added to the complex immediately after elution. Free receptor was separated from the complex by size exclusion chromatography on a Superdex 200 10/300 Increase column in 20mM Hepes pH 7.4, 100mM NaCl, 0,00075% MNG/2mol%, 0,00025% GDN, FLAG peptide, 5 mM EDTA, 1uM LM189, 0.1uM cmpd-6FA. Peak fractions were concentrated to 15-20mg/ml, filtered and used for electron microscopy experiments.

### Cryo-EM data collection and processing

3 μL aliquot of the β2AR-Gi-scfv16 complex was applied onto glow-discharged 300 mesh grids (Ultraufoil R1.2/1.3 or Quantifoil R1.2/1.3) and vitrified using a Vitrobot Mark IV (Thermo Fischer Scientific) under 100% humidity and 4°C conditions. Cryo-EM data were collected on a Titan Krios electron microscope operating at 300 kV and equipped with a K3 direct electron detector. Movies were acquired with a calibrated pixel size of 1.111 Å/pixel and a total dose of ∼52.5 electrons/Å², fractionated across 50 frames (Fig.S2, Table S2).

Data processing was performed using RELION 3.1.2 and cryoSPARC 3.3.2(*61*). Initially, motion correction was carried out on the movies using RELION’s built-in implementation, followed by Contrast transfer function (CTF) estimation using CTFFIND4(*62*). Reference-based particle picking utilized previously determined GPCR-G protein 2D classes. For the ultrafoil grid dataset, 3,659,953 particles were picked, subjected to 2D classification to remove low-quality particles, and further sorted through two rounds of 3D classification. This process yielded 265,559 particles and a 3.4 Å resolution structure after 3D refinement. In the case of the quantifoil grid dataset, 5,327,360 particles were picked, followed by one round of 2D classification and 3D classification, resulting in 513,747 particles and a 3.4 Å resolution structure. The two datasets were then merged, and a 3D classification without image alignment was performed, leaving 477,875 particles and a 3.3 Å resolution structure after 3D refinement.

Subsequent steps included CtfRefine, particle polishing, and the application of a mask to exclude the micelle and flexible alpha-helical domain. The final structure reached 2.9 Å resolution. The particles were then imported into cryoSPARC for non-uniform refinement, and the resulting map was sharpened using the Phenix autosharpen function to enhance map quality(*63*).

Lastly, local resolution estimation and 3DFSC were employed to assess the local resolution and orientation distribution of the final dataset(*64*) (Fig.S2).

### Model building and refinement

The individual structures of β_2_AR, Gi heterotrimer, and scfv16 were independently docked into the final sharpened map. Model and geometric restraints for LM189 and cmpd-6FA were generated using the Phenix elbow tool(*65*). Additionally, four cholesterol molecules were built into densities corresponding to previously identified cholesterol binding sites(*66*). The model was iteratively refined and validated through multiple rounds of Phenix real-space refinement and manual refinement in Coot (Table S2)(*67*).

### MD simulations

Simulations of the β_2_AR-Gi complex were based on the herein reported LM189-bound β_2_AR-Gi cryo-EM. The ligand LM189 was either kept in the orthosteric binding site or replaced by epinephrine by structurally aligning the cryo-EM with the epinephrine-bound β_2_AR-Nb6B9 structure (PDB-ID 4LDO)(*45*) and transferring the coordinates of epinephrine. Simulations of the β_2_AR-Gs complex were based on the BI-167107-bound β_2_AR-Gs crystal structure (PDB-ID 3SN6)(*4*). In order to obtain the LM189-bound and epinephrine-bound β_2_AR-Gs complexes, the coordinates of BI-167107 were removed and replaced by the coordinates of LM189 and epinephrine after structurally aligning the herein reported LM189-bound β_2_AR-Gi cryo-EM or the epinephrine-bound β_2_AR-Nb6B9 structure (PDB-ID 4LDO) onto the β_2_AR crystal structure, respectively. For further comparison, the salmeterol-bound β_2_AR-Nb71 structure (PDB-ID 6MXT) was subjected to MD simulations.

The five receptor complexes (LM189-β_2_AR-Gi, LM189-β_2_AR-Gs, epinephrine-β_2_AR-Gs, epinephrine-β_2_AR-Gi and salmeterol-β_2_AR-Nb71) were further prepared using UCSF Chimera(*68*). In order to save computational resources, we conducted all MD simulations without intracellular proteins but applied position restraints on all receptor residues within 5Å of the G protein or the Nb71 in order to maintain the respective conformation of the β_2_AR. The three missing amino acids 176-178 in the ECL2 of the β_2_AR-Gs crystal structure (PDB-ID 3SN6) were modeled by means of MODELLER(*69*), hydrogens were added, auxiliary proteins were removed and chain termini capped with neutral acetyl and methylamide groups. Except for Asp^2.50^ and Glu^3.41^, all titratable residues were left in their dominant protonation state at pH 7.0. Asp^2.50^ has been suggested to be protonated in the active state(*70*), and residue Glu^3.41^ directly contacts the lipid interface and therefore will also exist predominantly in its protonated state(*71*) (*72*). Thus, these residues were protonated in MD simulations. LM189, epinephrine and salmeterol were protonated at the secondary amine allowing the formation of the canonical salt bridge to Asp^3.32^ conserved in aminergic GPCRs.

Parameter topology and coordinate files of the four receptor complexes were build up using the leap module of AMBER18(*73*). Parameters for ligands were assigned using antechamber(*73*). Therefore, the structures of LM189, epinephrine and salmeterol were optimized by means of Gaussian16(*74*) at the B3LYP/6-31G(d) level (attributing a formal charge of +1), charges were calculated at the HF/6-31G(d) level and atom point charges assigned according to the RESP procedure(*75*). Energy minimization was performed applying 500 steps of steepest decent followed by 4500 steps of conjugate gradient. The protein structures were aligned to the Orientation of Proteins in Membranes (OPM)(*76*) Gs-bound structure of β_2_AR (PDB-ID 3SN6). Each complex was inserted into a pre-equilibrated membrane of dioleyl-phosphatidylcholine (DOPC) lipids by means of the GROMACS tool g_membed(*77*). Subsequently, water molecules were replaced by sodium and chloride ions to give a neutral system with 0.15 M NaCl. The final system dimensions were roughly 80 × 80 × 100 Å(*68*), containing about 156 lipids, 58 sodium ions, 67 chloride ions, and about 13,200 water molecules. For all simulations, the general AMBER force field(*78*) (GAFF2) was used for ligands, the lipid14 force field(*79*) for DOPC molecules, and ff14SB(*80*) for the protein residues. The SPC/E water model(*81*) was applied.

Simulations were performed using GROMACS 2021.1(*82–84*). The simulation systems were energy minimized and equilibrated in the NVT ensemble at 310 K for 1 ns followed by the NPT ensemble for 1 ns with harmonic restraints of 10.0 kcalꞏmol^-1^ on protein and ligands. In the NVT ensemble the V-rescale thermostat was used. In the NPT ensemble the Berendsen barostat, a surface tension of 22 dynꞏcm^-1^, and a compressibility of 4.5 × 10-5 bar^-1^ was applied. During the equilibration and the subsequent productive MD runs, position restraints of 10.0 kcalꞏmol^−1^were applied on the β_2_AR-residues within 5 Å of the G protein interface.

Multiple simulations were started from the final snapshot of the equilibration procedure for each of the four receptor complexes, resulting in productive molecular dynamics simulation runs of 3 × 2 µs for each simulation system. Simulations were performed using periodic boundary conditions and a time step of 2 fs with bonds involving hydrogen constrained using LINCS(*85*). Long-range electrostatic interactions were computed using particle mesh Ewald method(*86*) with interpolation of order 4 and FFT grid spacing of 1.6 Å. Non-bonded interactions were cut off at 12.0 Å. The analysis of the trajectories was performed using the CPPTRAJ module(*87*) of AMBER18. Interaction frequencies, distances and dihedrals were plotted using Matplotlib 2.2.2(*88*).

The equilibrated LM189-bound β_2_AR-Gi and epinephrine-bound β_2_AR-Gs complexes were further subjected to unrestrained MD simulations. In these simulations, following the same protocols outlined above, the position restraints on residues near the G protein interface were removed to allow for greater conformational flexibility. For the epinephrine-bound β_2_AR binary complex, 16 independent replicates were performed, with simulation times ranging from 500 ns to 6.4 μs. For the LM189-bound β_2_AR binary complex, 10 independent replicates were conducted, with simulation times ranging from 600 ns to 11 μs.

### Continuous-Wave EPR Spectroscopy

β2Δ5 with the Q142C mutation was expressed, purified and labeled as described above. Frozen SEC pure receptor aliquots in 20 mM HEPES pH 7.4, 100 mM NaCl, 0.01% LMNG were thawed and incubated with ligands at 10-X molar excess for 1 hour at room temperature; buffer matched or G protein was added at 2X molar excess to aliquots after 1 hour ligand incubation and incubated for another 2 hours. Apyrase (1 unit, NEB) was added for an additional 1.5 hours. Samples were then loaded into a quartz capillary (0.9 mm ID, 1.3 mm OD; #2-000-050, Drummond Scientific Company) with a volume of approximately 30 μL.CWEPR spectroscopy was performed at X-band (∼9.46 GHz) on a Bruker Magnettech ESR5000 spectrometer at room temperature. Spectra were recorded at a microwave power of 36 mW with 100 KHz field modulation at a modulation amplitude of 0.1 mT, scan width of 20 mT, and a scan rate of 0.24 mT/s. CW data were aligned and baseline corrected using the custom software programs Convert&Align101 and Baseline048 written in LabVIEW by Dr. Christian Altenbach (University of California, Los Angeles) and freely available upon request. Processed CW-EPR data were plotted in Graphpad Prism 9.3.1.

### DEER Spectroscopy

DEER samples consisted in approximately 100 µM spin-labeled β2Δ6 N148C/L266C, 10-X molar excess of ligand, 2x molar excess of G proteins or matched deuterated buffer. After 2 hours incubation with the G protein, apyrase (1 unit NEB) was added for an additional 1.5 hours. Finally, glycerol-d_8_ was added as a cryoprotectant to a final concentration of 20% (v/v). Samples were loaded into borosilicate capillaries (1.4 mm ID, 1.7mm OD; VitroCom) at final volumes of 14-20 μL and flash frozen in liquid nitrogen.

Experiments were performed at Q-band (∼33.65 GHz) on a Bruker Elexsys E580 equipped with a SpinJet AWG, EN5107D2 resonator, variable-temperature cryogen-free cooling system (ColdEdge Technologies Inc.), and either a 10 W solid state amplifier or a 150 W TWT amplifier (Applied Systems Engineering Inc.). The 150 W TWT amplifier was used to improve signal-to-noise with respect to modulation for the transducer-coupled samples; data for all other samples were collected using the 10 W amplifier. Data were collected at a temperature of 50 K.

Dipolar evolution data were acquired using the dead-time free 4-pulse DEER sequence (π/2)_obs_ – (d_1_) – (π)_obs_ – (d_1_ + T) – (π)_pump_ – (d_2_ - T) – (π)_obs_ – (d_2_) – echo(*89*),(*90*) with 16-step phase cycling(*91*). The parameters used for DEER experiments are listed in Table S3. Gaussian shapes were implemented for all pulses using the built-in Gaussian pulse profile in Bruker Xepr software(*92*); resonator bandwidth compensation was not used. Pump pulses were applied to the maximal intensity of the field swept echo detected absorption spectrum. Observe pulses were applied at a frequency either 45 MHz or 90 MHz lower than the pump pulses for experiments performed with the 10 W or 150 W amplifier, respectively. Optimal microwave power for π/2 and π pulses was determined by adjusting pulse amplitudes for a transient nutation experiment to achieve maximum Hahn echo inversion at the pulse lengths being used (72 ns)(*93*).

DEER data were processed with ComparativeDeerAnalyzer (CDA) automated processing in DeerAnalysis2021(*94*). This utilizes two different fitting routines: neural network analysis with DEERNet(*95*) from Spinach revision 5501 and Tikhonov regularization with DeerLab 0.9.1 routines(*96*). The resulting consensus fit is a mean of the two with a 95% confidence interval reported that is comprised of both method’s errors. DEER time traces and distance distributions for all samples are shown in Fig.S5E; time traces are normalized to the signal intensity at time = 0 and distance distributions are area-normalized. DEER data were plotted in Graphpad Prism 9.3.1.

### smFRET Spectroscopy

#### smPull receptor isolation and surface display

To inhibit nonspecific protein adsorption, flow cells for single-molecule experiments were prepared as previously described(*50*) using mPEG (Laysan Bio) passivated glass coverslips (VWR) and doped with biotin PEG16. Before each experiment, coverslips were incubated with NeutrAvidin (Thermo), followed by 10 nM biotinylated antibody (mouse anti-FLAG, Jackson ImmunoResearch). Between each conjugation step, the chambers were flushed to remove free reagents. The antibody dilutions and washes were done in T50 buffer (50 mM NaCl, 10 mM Tris, pH 7.5). To achieve sparse immobilization of labeled receptors on the surface, purified labeled receptor was diluted (ranging from 100X to 1000X dilution) and applied to coverslips. After achieving optimum surface immobilization (∼400 molecules in a 2,000 μm^2^ imaging area), unbound receptors were washed out of the flow chamber and the flow cells were then washed extensively (up to 50X the cell volume).

#### smFRET measurements

Receptors were imaged for smFRET in imaging buffer consisting of (in mM) 3 Trolox, 100 NaCl, 2 CaCl_2_, 20 HEPES, 0.01% MNG and an oxygen scavenging system (0.8% dextrose, 0.8 mg ml^-1^glucose oxidase, and 0.02 mg ml^-1^ catalase), pH 7.4. All buffers were made in UltraPure distilled water (Invitrogen). Samples were imaged with a 1.65 na X60 objective (Olympus) on a total internal reflection fluorescence microscope with 100 ms time resolution unless stated otherwise. Lasers at 532 nm (Cobolt) and 633 nm (Melles Griot) were used for donor and acceptor excitation, respectively. Fluorescence passed through Chroma ET550lp and split into donor and acceptor signal with a Chroma T635lpxr. FRET efficiency was calculated as (*I*_A_-0.1*I*_D_)/(*I*_D_+*I*_A_), in which *I*_D_ and *I*_A_ are the donor and acceptor intensity, respectively, after back-ground subtraction. Movies were recorded at 100 ms acquisition time (10 Hz) with a Photometrics Prime 95B cMOS camera using micromanager acquisition software.

#### smFRET data analysis

SPARTAN version 3.7(*97*) was used to analyze fluorescent movies. Donor and acceptor channels were aligned using the first 10 frames of each movie while excluding particles closer than 3.5 pixels and using an integration window of 12 pixels. Single-molecule intensity traces showing single-donor and single-acceptor photobleaching with a stable total intensity for longer than 5s (50 frames), SNRbg > 15 and donor acceptor correlation coefficient < 0.0 were collected (20–30% of total molecules per imaging area). Individual traces were smoothed using a nonlinear filter(*98*) with following filter parameters: window = 2, M = 2 and P = 15. Each experiment was performed ≥4 times to ensure reproducibility. smFRET histograms were compiled from ≥100 molecules per condition (100 ms time resolution). Error bars in the histograms represent the standard error from ≥4 independent movies. To ensure that traces of different lengths contribute equally, histograms from individual traces were normalized to one before compiling. Gaussian fitting to histograms was done in Origin Pro.

#### Fluorescence spectroscopy

Fluorescence experiments were performed on a Fluoromax 4C spectrofluorometer (Horiba Scientific, Edison, NJ, USA) using 5 nm excitation slit width and 3nm emission slit width. Emission spectra were recorded using excitation at 380 nm. Concentrations after mixing were as follows: β_2_AR – 100 nM, salmeterol – 100 µM, LM189 – 75 µM, BI-167107 – 25 µM. Ligand concentrations were chosen to achieve receptor saturation. Experiments were conducted in buffer containing 20mM Hepes pH 7.4, 100mM NaCl, 0.01% LMNG. Samples were measured after one hour incubation in the dark at the final concentrations to allow full equilibration. Measurements were performed in biological triplicates.

#### Gi and Gs coupling in intact HEK cells

HEK-A cells (or HEK-ΔGNAS) were co-transfected with rLuc-tagged β_2_AR (β_2_AR-rLuc8) and the Venus-miniGs (venus mGs) sensor containing the C-terminal residues from either Gα_s_, or Gα_i1_ (*11*, *99*, *100*) and a nuclear export signal (NES-venus-mGs) and allowed to express for 48 h at 37C in DMEM in a CO_2_ incubator. These chimeras will be referred to as mGs and mGs/i. Expression vectors containing β_2_AR-rLuc8 and the mGs chimeras were generously provided by Nevin Lambert (Augusta University at Georgia). Transfected cells were harvested with EDTA (2 mM) in PBS, washed (by centrifugation) in a Hepes buffered Saline Solution (HBSS), and resuspended in HBSS containing 0.1% ascorbic acid and 1% DMSO. Cells were transferred into 96-well plates (100 mL per well) and incubated for 20 min at RT. Cells were then preincubated with coelentrazine (5 mM final) for 5 minutes prior to the addition of agonist. An agonist dose-response relationship was performed through the addition of a 10X agonist concentration to the cells, and the fluorescence emission measured (at 485 and 530 nm) using a Molecular Devices M5 fluorescence plate reader. The data were collected in kinetic mode every 120 s for 30 min total. Activity values were derived from the area under the BRET ratio (em530/em485) progress curve between 6 and 16 min. Data were analyzed using Prism (GraphPad™, La Jolla CA).

## Acknowledgements

We thank Elizabeth Montabana for assistance with the Cryo-EM data collection. We acknowledge Alem W. Kahsai for cmpd-6FA. We acknowledge National Institutes of Health Instrumentation grant S10OD024980 in support of acquisition of the UC Santa Cruz pulsed EPR spectrometer. We thank Kayo Sato, Shigeko Nakano and Ayumi Inoue (Tohoku University) for their assistance in the NanoBiT assay. We thank Dr. Nevin Lambert (Augusta University) for the β_2_AR-rLuc8 and miniG constructs as well as helpful discussion. Cryo-EM data were collected at Stanford Cryo-Electron Microscopy Center (cEMc).

The materials used in this study can be provided by BKK, MTL and PG pending scientific review and a completed material transfer agreement. Requests for the materials should be submitted to: MC, BKK, MTL and PG.

## Funding

M.C. received funding from the European Union Horizon 2020 research and innovation program under the Marie Sklodowska-Curie grant agreement n° 799376.

M.C. was supported by the American Heart Association (AHA) Postdoctoral Fellowship award number 915188.

B.K.K. was supported by R01NS028471.

M.T.L. is supported by R01GM135581 and grant S10 OD025260.

P.G. is supported by the Deutsche Forschungsgemeinschaft (DFG, German Research Foundation) DFG grants GRK 1910 and GM 13/14-1.

R.J.L. is an investigator with the Howard Hughes Medical Institute and is supported by R01HL0160371609 from the National Heart, Lung, and Blood Institute, NIH.

P.B is supported by F31 HL164002.

E.Y.I. is a Weill Neurohub Investigator and is supported by R01NS119826.

N.N. was supported by NSF MCB-1942957, American Chemical Society Petroleum Research Fund, 61678-UR6.

Y.K.X. is supported by NIH grant HL162825 and MH134119 and VA Merit grants IK6BX005753 and I01BX005100.

A.I. is funded by the Society for the Promotion of Science (JP21H04791 and JP24K2128); the Japan Science and Technology Agency (JPMJFR215T and JPMJMS2023); the Japan Agency for Medical Research and Development (JP22ama121038 and JP22zf0127007).

R.K.S is funded by National Institute of General Medical Sciences grants GM083118.

## Author contributions

Conceptualization: MC, BKK, MTL, HW, RJL, BP, PB

Methodology: MC, BKK, MTL, PG, RJL, NN, EYI, AI, RKS, HW, BP, PB, LM, SMFM

Software: HW

Validation: MC, BKK, PG, PB, RKS, NN, SMFM

Formal analysis: MC, PB, RKS, NN, CH, LM, SMFM, YKX

Investigation: MC, HW, PB, NN, HH, LM, MFS, SMFM, BX, RKS, TEA, EW, AI, TEA, NS, AI, BC

Resources: BKK, PG, MTL, MC, BP, PB, LM, YKX, HW

Data Curation: NN, HW

Visualization: MC, HW, PB, CH, HH, MFS, AI, NS, RJL, EYI, NN, SMFM, YKX

Funding acquisition: MC, BKK, MTL, PB, PG, RJL, EYI, NN, YKX, AI, RKS

Project administration: MC, BKK, MTL, PG, RJL

Supervision: BKK, PG, MTL, MC, RJL, EYI, YKX

Writing – original draft: MC, BKK, HW, PB

Writing – review & editing: MC, BKK, MTL, PG, RJL, NN, EYI, PB, BP, AI, NS, LM, MFS, AI, BC, SMFM

## Competing interests

B.K.K. is a co-founder of and consultant for ConfometRx, Inc. R.J.L. is a founder of Trevena, Inc. and a co-founder and stockholder of Septerna. R.J.L. is also a co-founder and stockholder of Lexicon Pharmaceuticals. B.P. is a consultant with Septerna. The other authors declare no competing interests.

## Data and materials availability

The cryo-EM density map has been deposited in the Electron Microscopy Data Bank (EMDB) under accession code EMD-44925, and model coordinates have been deposited in the Protein Data Bank (PDB) under accession ID 9BUY. Materials described in this study are available upon request sent to the corresponding authors. All data needed to evaluate the conclusions in the paper are present in the paper and/or the Supplementary Materials.

## Notes

### Competing Interest Statement

B.K.K. is a co-founder of and consultant for ConfometRx, Inc. R.J.L. is a founder of Trevena, Inc. and Septerna. R.J.L. is also on the board of Lexicon Pharmaceuticals. B.P. is a consultant with Septerna.

### Summary of Updates

Some experimental data has been removed: nano-BiT recruitment assays, cardiomyocyte activity assays. Figure 1 and Supplemental files updated.

