## Supplemental Information for "Structure and dynamics determine G protein coupling specificity at a class A GPCR Teaser: Structure and dynamics studies reveal a mechanism for GPCR signaling bias and G-protein coupling specificity"

**The PDF file includes:**

Fig. S1 to S6
Tables S1 to S3

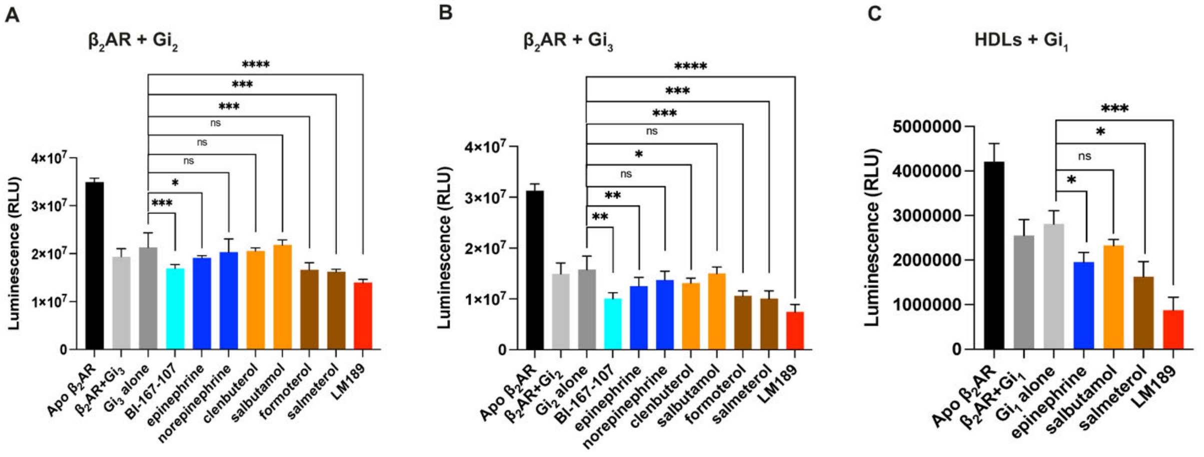


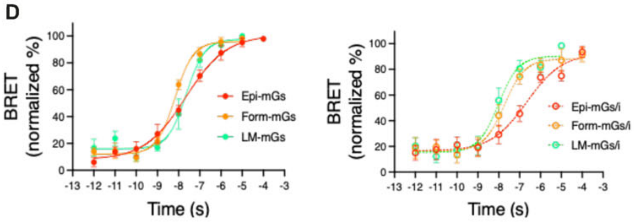


| | Gs | | |
|---|---|---|---|
| Ligand | Log(t/KA) | ΔLog(t/KA) | RE |
| Epinephrine | 7.66±0.17 | 0 | 1 |
| Formoterol | 8.19±0.08 | 0.5±0.19 | 3.16* |
| LM189 | 7.63±0.14 | -0.03±0.22 | 0.93* |

*=*P*<0.05

| | Gi | | |
|---|---|---|---|
| Ligand | Log(t/KA) | ΔLog(t/KA) | RE |
| Epinephrine | 6.65±0.31 | 0 | 1 |
| Formoterol | 7.72±0.19 | 1.07±0.36 | 11.72* |
| LM189 | 8.01±0.18 | 1.35±0.35 | 22.39* |

*=*P*<0.05

| Ligand | ΔΔLog(t/KA) | BF |
|---|---|---|
| Epinephrine | 0 | 1 |
| Formoterol | 0.57±0.4 | 3.7* |
| LM189 | 1.38±0.41 | 24* |

*=*P*<0.05

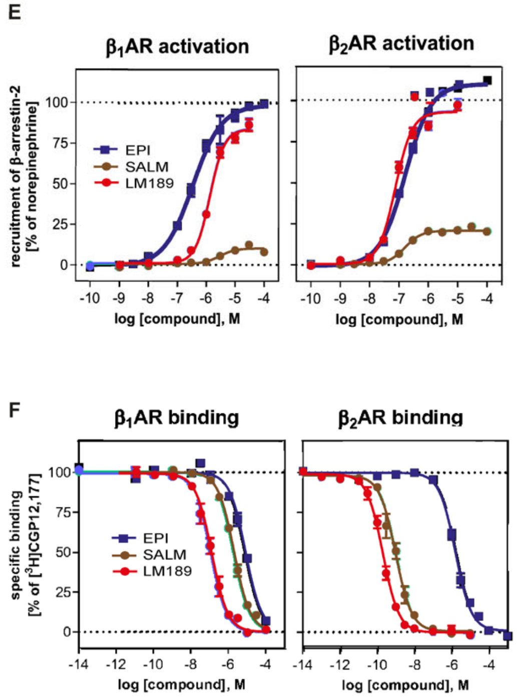


**Fig. S1.**

**Characterization of the LM189 Gi-biased ligand. (A-B)** Luminescence GTP-ase-Glo assays of $\beta_2$AR coupled to (**A**) G$\alpha i_2$ and (**B**) G$\alpha i_3$ in the presence of ligands with different efficacies. (**A**) Salmeterol promotes higher G$\alpha i_2$ turnover compared to the G-protein alone control (***P=0.0004). LM189 is more efficacious than salmeterol (****P≤0.0001). **(B)** Salmeterol promotes higher Gi$_3$ turnover compared to the G-protein alone control (**P=0.0019). LM189 is more efficacious than salmeterol (*****P*<0.0001) **(C)** GTP-ase-Glo assay of $\beta_2$AR reconstituted in HDLs particles and coupled to G$\alpha i_1$ in the presence of ligands with different efficacies. LM189 promotes higher G-protein turnover compared to G$\alpha i_1$ alone (***P=0.0003). In A-C, data are presented as the mean ± s.d. of biological triplicates. P values were calculated using the unpaired t test analysis on GraphPad Prism, assuming Gaussian distributions. Ns=(*P*>0.05), * (*P*≤0.05), ** (*P*≤0.01), *** (*P*≤0.001), **** (*P*≤0.0001). **(D)** LM189 bias quantification. Top panels: concentration-response curves of Gs (left) and Gi (right) activation mediated by epinephrine (red), formoterol (orange) and LM189 (green). Data points represent the mean ± s.d. of n=5 experiments. Upper and middle table: Δlog(τ/KA) ratios for Gs (upper table) and Gi (middle table) calculated from the log(τ/KA) ratios obtained from dose-response curves, considering epinephrine as the reference ligand. The relative effectiveness (RE) of the ligands toward each pathway, relative to epinephrine, was determined. Lower table: $\Delta\Delta\log(\tau/KA)$ ratios calculated from the $\Delta\log(\tau/KA)$ ratios. The ligand bias factors (BF), relative to epinephrine, were determined. Data were analyzed in a pairwise manner using a two-tailed unpaired student's t-test to determine the significance of the ligand biases, where $P<0.05$ was considered to be significant. **(E)** β-Arrestin-2 recruitment of epinephrine-bound and LM189-bound $\beta_1$- and $\beta_2$AR. Data are represented as the mean ± s.d. of 6-17 independent experiments each conducted in duplicates. **(F)** Competition binding experiments of LM189 at the $\beta_1$- and $\beta_2$-adrenoreceptor in comparison to the reference ligands epinephrine and salmeterol. Data are represented as the mean ± s.d. of 7-11 single experiments each performed in triplicates.

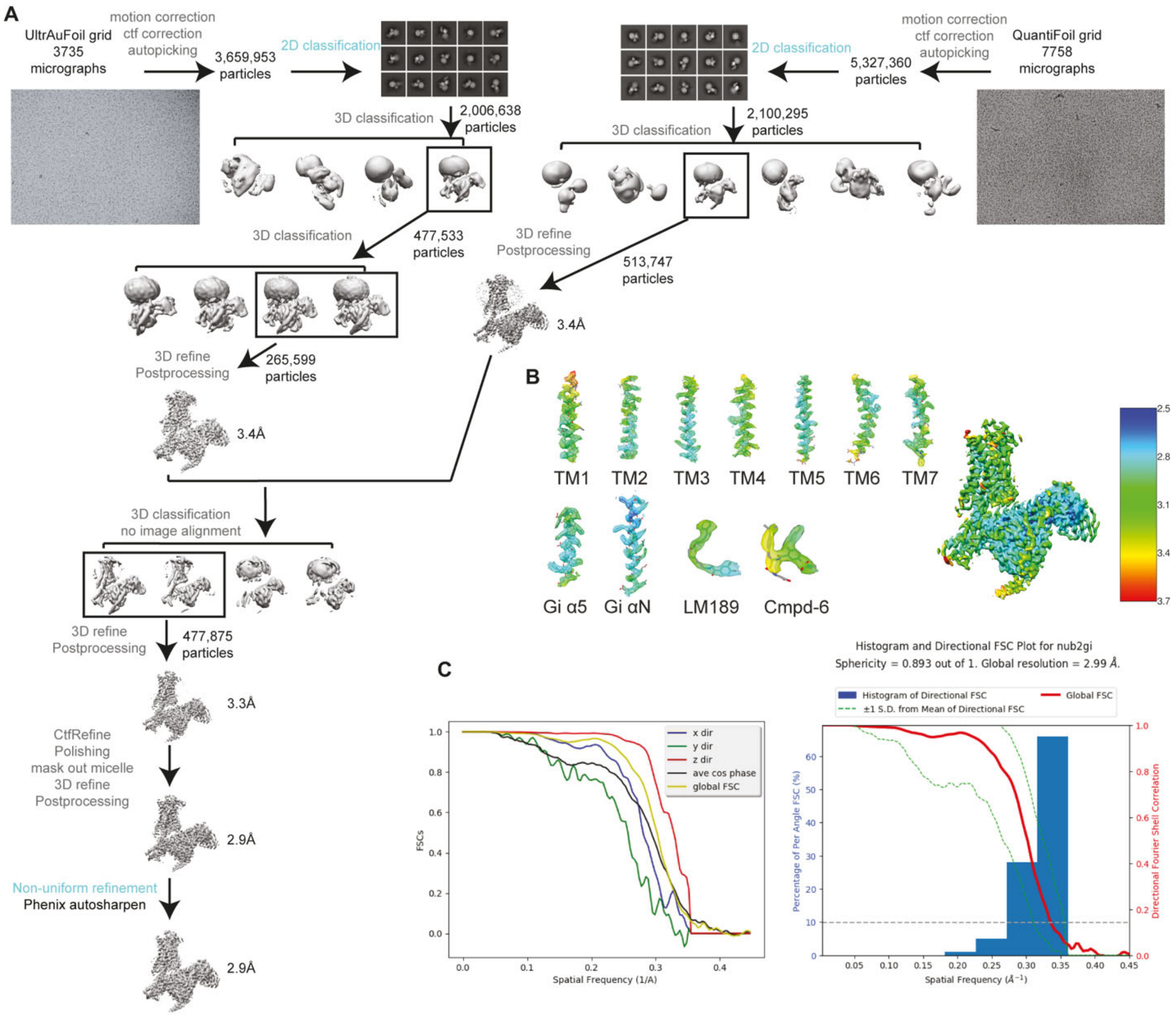


**Fig. S2.**

**Cryo-EM data processing workflow and validation (A)** cryo-EM data processing workflow. For detailed information please refer to Materials and Methods section. **(B)** Local resolution information of the final cryo-EM map. Local resolution colored EM densities and models are also shown for TM1-TM7, Gi α5, Gi αN, LM189, and Cmpd-6FA. Densities were visualized with UCSF ChimeraX (*68*). **(C)** Directional resolution validated by 3DFSC.

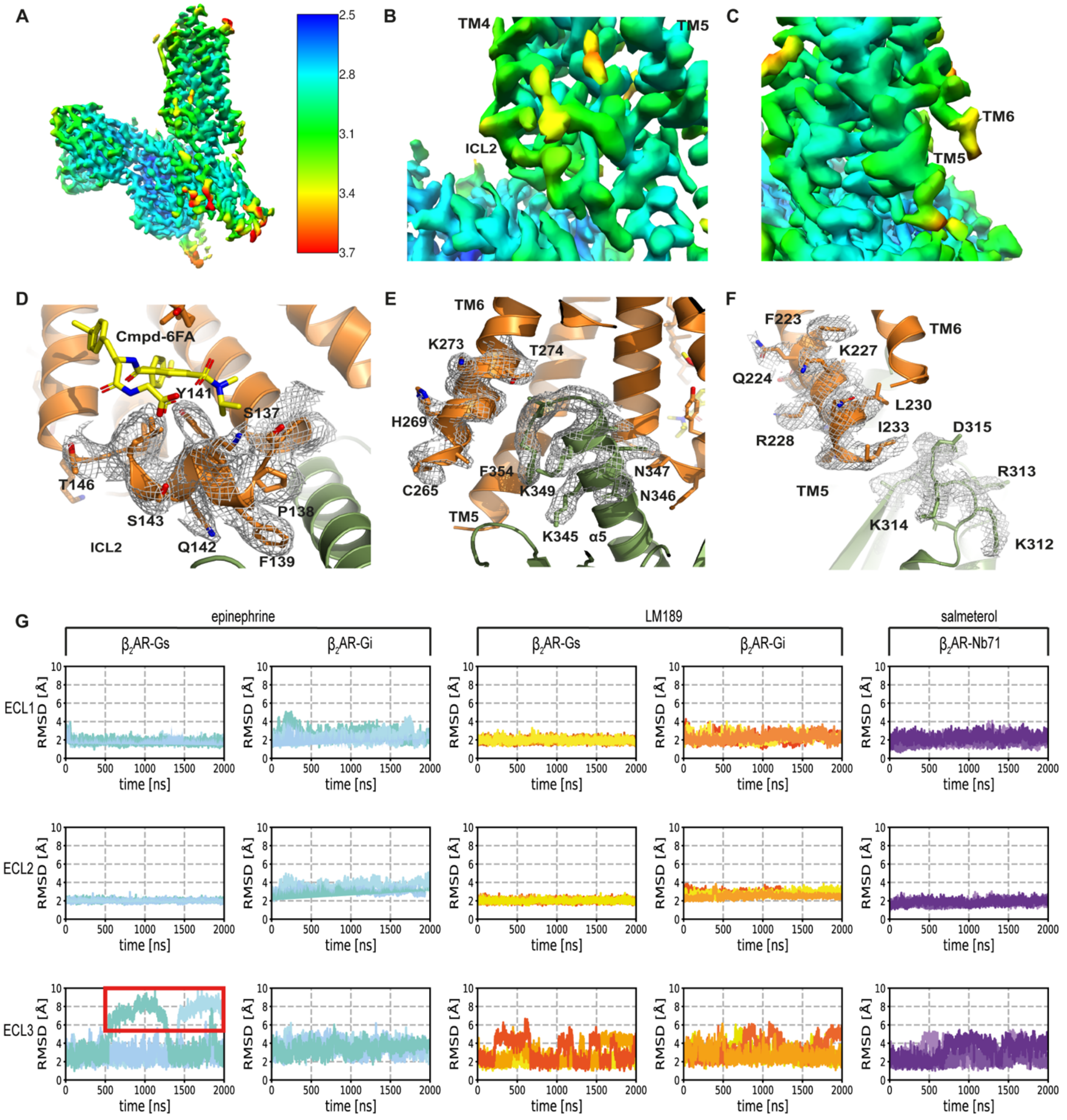


### Fig. S3.

**The LM189-bound β$_2$AR-Gi complex structure.** **(A-C)** Local resolution Cryo-EM estimate maps. **(A)** Local resolution estimate map of the LM189-bound β$_2$AR-Gi complex. **(B)** Close-up of the local resolution estimate map around ICL2 of β$_2$AR. **(C)** Close-up of the local resolution estimate map around TM5 of β$_2$AR. **(D-F)** Cryo-EM densities of the β$_2$AR-Gi complex **(D)** Density around ICL2 of the β$_2$AR-Gi complex **(E)** Density at the tip of TM6 and the alpha5 of Gi. **(F)** Density around the tip of TM5 and the GTP-ase domain of Gi. **(G)** MD simulations of the extracellular region of ligand-bound β$_2$AR active state. From left to right, replicates of epinephrine-bound β$_2$AR-Gs and β$_2$AR-Gi simulations are showed in aquamarine, light-blue and turquoise, LM189-bound β$_2$AR-Gs and β$_2$AR-Gi replicates are colored in orange, red and yellow and salmeterol-bound β$_2$AR-Nb71 replicates are colored in violet and lilac.

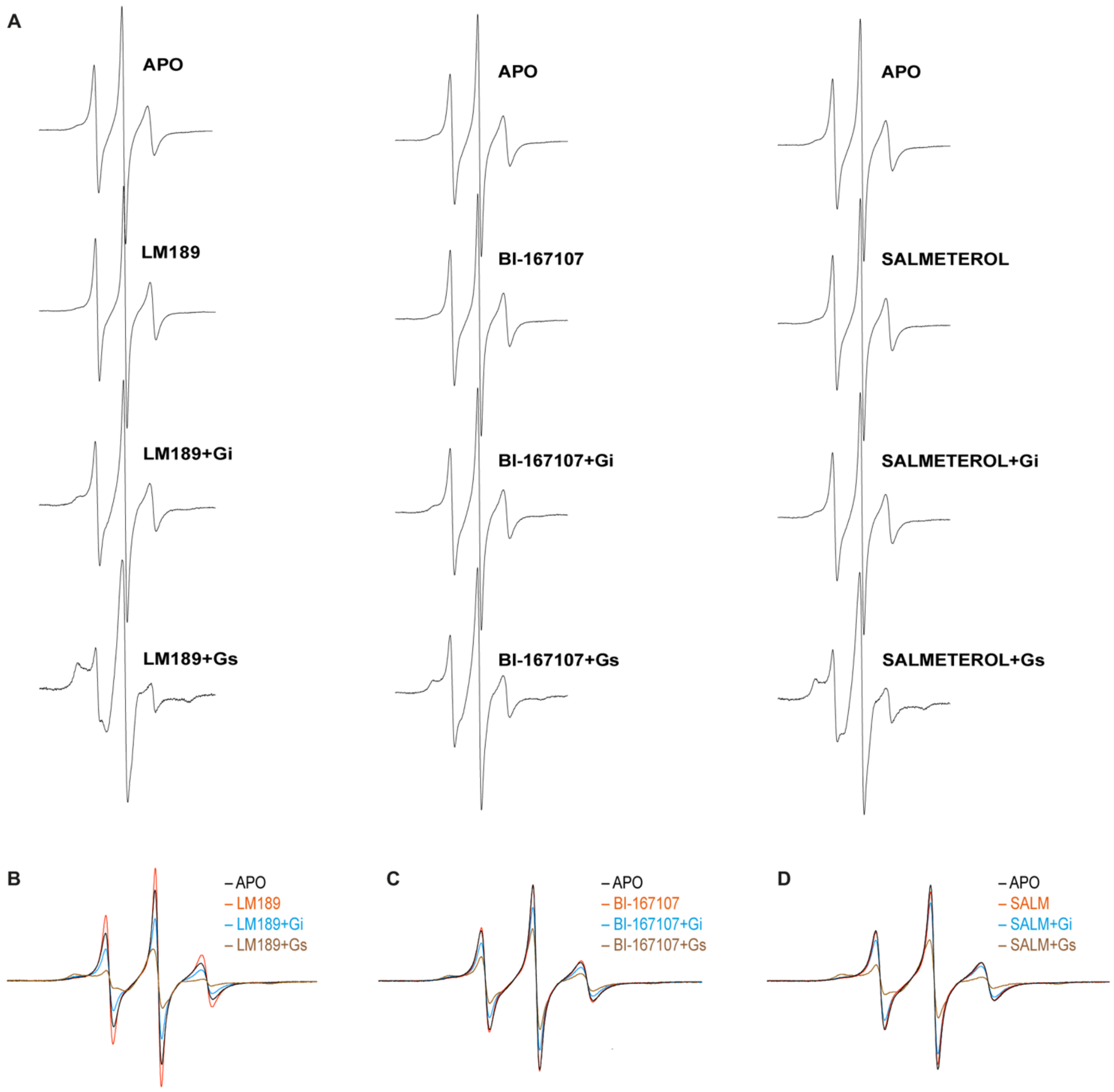


#### Fig. S4.

**Cw-EPR studies of β₂AR ICL2 dynamics. (A)** Amplitude normalized 142-IAP spectra for each of the experimental conditions tested. **(B-D)** Area-normalized 142-IAP spectra of the apo, ligand-bound, and G protein-bound receptor. Superimposed spectra are shown for the apo and **(B)** LM189, **(C)** BI-167107, and **(D)** salmeterol conditions.

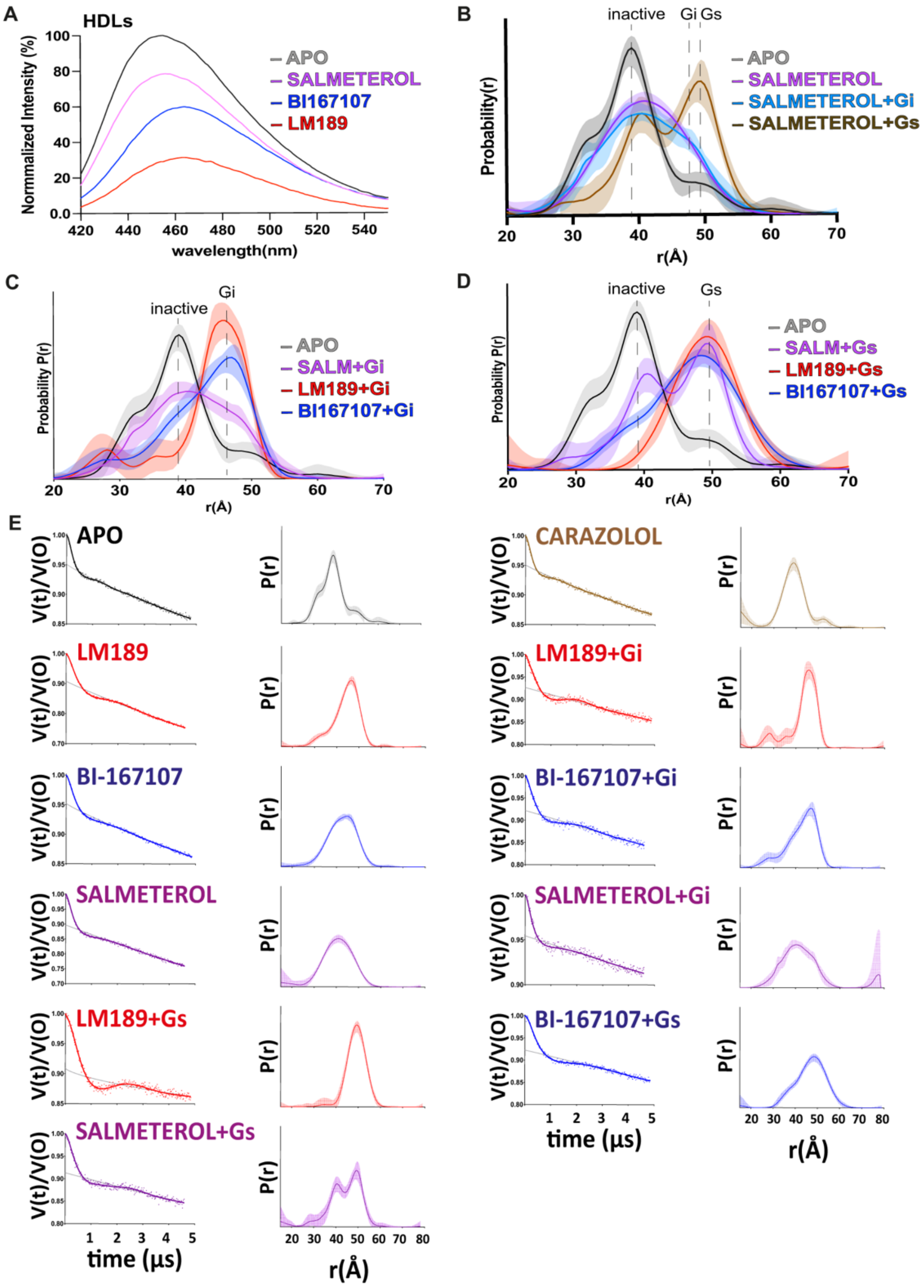


#### Fig. S5.

**Investigation of TM6 conformational dynamics. (A)** Steady-state fluorescence emission spectra of mBBr-labeled β$_2$AR reconstituted in HDLs, in the presence of ligands. The spectra are normalized relative to apo (unliganded) receptor (gray). **(B-E)** DEER investigations of TM4/6 of β$_2$AR in LMNG/CHS. Distance distributions in the presence of **(B)** salmeterol and G proteins, **(C)** Gi-bound receptor in the presence of different ligands, and **(D)** Gs-bound receptor in the presence of different ligands. **(E)** Background-subtracted dipolar evolution functions and distance distributions are shown for each sample. All data are color-coded as indicated.

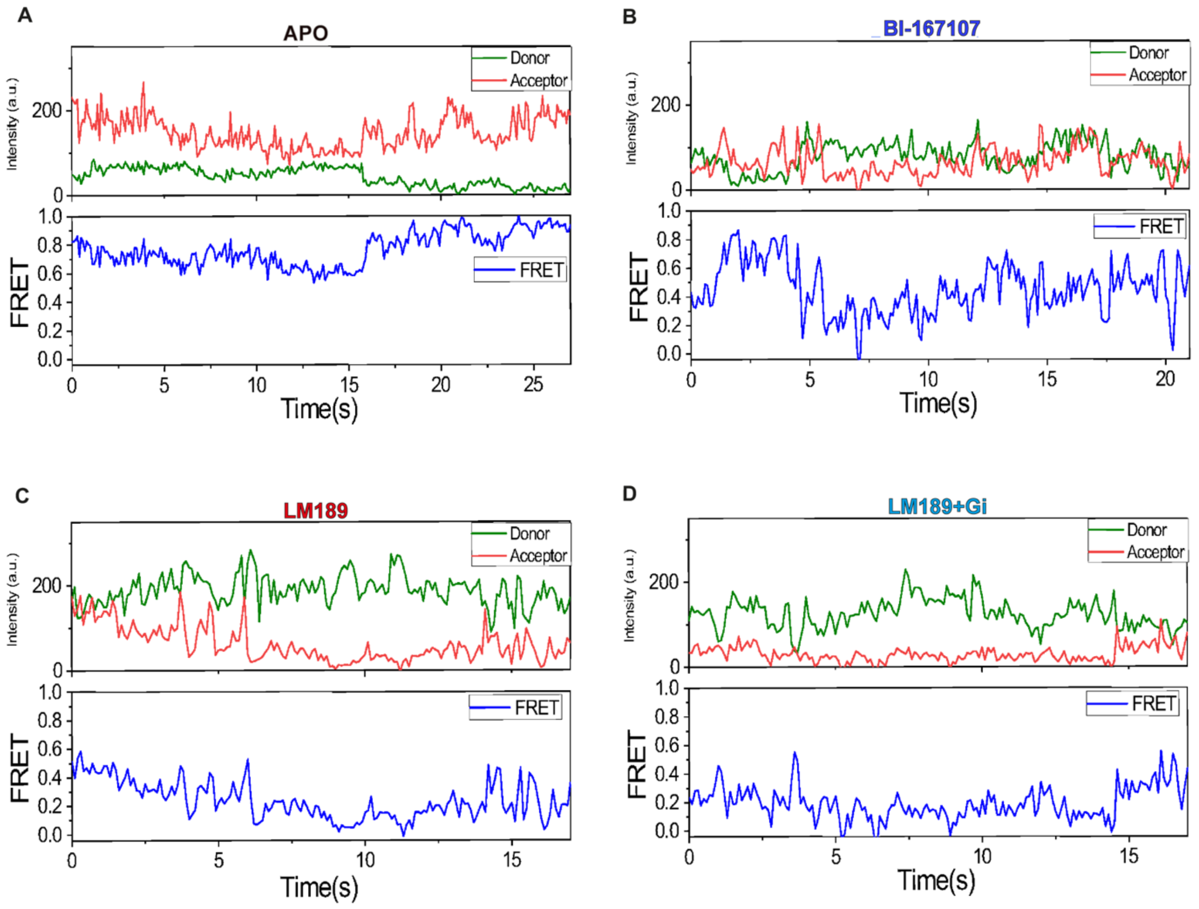


**Fig. S6.**

**SmFRET representative traces.** Donor and acceptor intensity values are reported in green and red respectively. Calculated FRET values are shown in blue. Example traces and FRET values are displayed for the apo condition **(A)**, for the full agonist BI-167107 **(B)**, for the Gi-biased ligand LM189 **(C)** and for LM189-bound receptor coupled to Gi and scfv-16 **(D)**.

| Receptor Binding | | | | |
|---|---|---|---|---|
| Compound | $\beta_1$AR | | $\beta_2$AR | |
| | Ki [nM ± S.E.M][b] | n[c] | Ki [nM ± S.E.M][b] | n[c] |
| LM189 | 33 ± 5.6 | 7 | 0.058 ± 0.009 | 10 |
| Epinephrine | 2200 ± 290 | 8 | 380 ± 33 | 11 |
| Salmeterol | 520 ± 47 | 10 | 0.31 ± 0.064 | 9 |

| β-arrestin 2 recruitment | | | | | | |
|---|---|---|---|---|---|---|
| Compound | $\beta_1$AR | | | $\beta_2$AR | | |
| | $EC_{50}$[nM ±S.E.M][e] | $E_{max}$[nM ±S.E.M][f] | n[g] | $EC_{50}$[nM ±S.E.M][e] | $E_{max}$[nM ±S.E.M][f] | n[g] |
| LM189 | 1300 ± 31 | 82 ± 3 | 6 | 71 ± 51 | 92 ± 3 | 6 |
| Epinephrine | 380 ± 52 | 100 ± 2 | 6 | 190 ± 6.9 | 111 ± 1 | 17 |
| Salmeterol | 3000 ± 820 | 10 ± 1 | 7 | 190 ± 16 | 21 ± 1 | 13 |

[a] Binding affinity determined in a radioligand displacement assay using membranes from HEK293T cells transiently transfected with the cDNA of $\beta_1$AR or $\beta_2$AR and the radioligand [$^3$H]CGP12,177. [b] Mean $K_i$ values in [nM ± S.E.M.]. [c] Number of individual experiments each performed in triplicates. [d] Receptor mediated recruitment of β-arrestin-2 measured in HEK293T cells stably expressing the enzyme acceptor (EA) tagged β-arrestin-2 fusion protein and transiently transfected with the $\beta_1$AR or $\beta_2$AR each fused to the ProLink-PK1 fragment for enzyme complementation. [e] Potency displayed as mean $EC_{50}$ value in [nM ± S.E.M.]. [f] Intrinsic activity $E_{max}$ in [% ± S.E.M.] relative to the maximum effect of norepinephrine. [g] Number of individual experiments each performed in duplicates

#### Table S1.

Receptor binding data and functional data for β-arrestin-2 recruitment of LM189 at the $\beta_1$- and $\beta_2$-adrenoreceptor in comparison to the references epinephrine and salmeterol.

| **Data collection and processing** | |
|---|---|
| Magnification | 82,000 |
| Voltage (kV) | 300 |
| Electron exposure | 52.5 |
| Defocus range (*μ*m) | 0.7-2 |
| Pixel size (Å) | 1.111 |
| Symmetry imposed | C1 |
| Initial particle images (no.) | 8987313 |
| Final particle images (no.) | 477875 |
| Map resolution (Å) | 2.9 |
| FCS threshold | 2.5-3.7 |
| Map resolution range (Å) | 2.5-3.7 |

| **Refinement** | |
|---|---|
| Initial model used (PDB code) | 3SN6, 6DDE |
| Model resolution (Å) | 3.2 |
| FCS threshold | 0.5 |
| Model resolution range (Å) | 3.2-28.9 |
| Map sharpening *B* factor ($Å^2$) | -83.99 |
| Model composition | |
| Non-hydrogen atoms | 8640 |
| Protein residues | 1117 |
| Ligands | 5 |
| *B* factors ($Å^2$) | 2.5-3.7 |
| Protein | 63.61 |
| Ligand | 74.5 |
| R.m.s. deviations | |
| Bond lenghts (Å) | 0.003 |
| Bond angles (°) | 0.59 |
| Validation | |
| MolProbity score | 1.72 |
| Clashscore | 7.65 |
| Poor rotamers (%) | 0 |
| Ramachandran plot | |
| Favored (%) | 95.6 |
| Allowed (%) | 4.37 |
| Disallowed (%) | 0 |

#### Table S2.

**Cryo-EM acquisition parameters of LM189-bound $\beta_2$AR-Gi$_1$-scFv16**

| Parameter | 10W solid state amp | 150W TWT amp |
|---|---|---|
| $\pi/2_{obs}$, $\pi_{obs}$, and $\pi_{pump}$ | 30.6 ns FWHM (72 ns time base) | 15.3 ns FWHM (36 ns time base) |
| $\Delta v$ (MHz) | 45 | 90 |
| $d_1$ (ns) | 400 | 400 |
| $d_2$ (ns) | 3000-5000 | 3000-5000 |
| Shot repetition time (μs) | 4000 | 4000 |
| Shots per point | 10 | 4 |
| Integration window (ns) | 72 | 36 |

#### Table S3.

**DEER data acquisition parameters.**

**Titles of excel tables in auxiliary materials:**

Table 1: dose-response BRET
Table 2 : bias quantification
